# Measuring neuronal input-output operations *in vivo* with random access projection microscopy

**DOI:** 10.64898/2026.11.05.764682

**Authors:** Michael E. Xie, Maedeh Seyedolmohadesin, Manni He, Lillian Li, Yukun Alex Hao, Daniel Flickinger, Ryan Gibiser, Jonathan King, Kort Driessen, Jimin Park, R. Carter Peene, Henry Loeffler, Katrina Nguyen, Madeline Tom, Ali Williford, Jackie Swapp, Giulio Tononi, Chiara Cirelli, Michael Z. Lin, Jérôme Lecoq, Bruce E. Kimmel, Jack Glaser, Matthew D. Golub, Adam S. Charles, Karel Svoboda, Georg Jaindl, Kaspar Podgorski

## Abstract

Neurons transform thousands of synaptic inputs into spiking output, and understanding this transformation requires measuring many submicron synapses at millisecond timescales *in vivo*. Here we present SLAP2, a two-photon random access projection microscope with no per-target access time cost, yielding over 5-fold faster imaging than existing methods, and performant real-time hardware >800 synapses volumetrically at 50–100 Hz, and compartment-resolved dendritic voltage at 2–10 kHz motion correction. Flexible scanning modes enable measurement of hundreds of synapses at `~`200 Hz, in behaving mice. Simultaneous dendritic glutamate and somatic calcium imaging in visual cortex pyramidal neurons revealed nonlinear input-output amplification of stimulus onset and preferred stimulus orientation responses. Amplified features are associated with spatially organized inputs and stronger trial-to-trial coupling between synaptic activity and somatic output. These measurements establish SLAP2 as a powerful tool for recording intricate patterns of subcellular fluorescence dynamics in behaving animals.

## Introduction

The mammalian brain contains millions to billions of interconnected neurons that mediate cognition and behavior. Each neuron performs computations by combining thousands of synaptic inputs from other neurons to produce action potential output. Artificial neural networks based on this principle, with interconnected nodes each performing a simple nonlinear transformation, can compute any well-behaved function with arbitrary precision (Hornik et al. 1989). A network’s computation is specified by the inputs each neuron receives, their weights, and the nonlinear function it applies. Measuring the analogous quantities in the living brain, such as synaptic input patterns to individual neurons and how they are transformed into spiking output, is a critical step toward understanding neural computation.

Fluorescent indicators for glutamate allow imaging of activity at individual synapses with high sensitivity (Aggarwal et al. 2026). Similarly, recent advances in voltage indicators enable sensitive recordings of subthreshold membrane potential changes and spikes (Hao et al. 2026; Hao et al. 2024; Villette et al. 2024; Abdelfattah et al. 2023; Liu et al. 2022). However, recording these indicators across a representative number of synapses and dendritic compartments with sufficient speed to study computations in the intact brain remains a challenge.

One difficulty is the minuscule size of the structures involved. Synapses have sub-femtoliter volumes and are often separated by less than one micrometer in grey matter (Parajuli et al. 2020; DeFelipe et al. 1999). These spatial scales require two-photon microscopy, which retains high resolution and contrast deep in tissue (Svoboda & Yasuda 2006; Denk & Svoboda 1997). Neurons receive synapses on tree-like dendrites, which fill less than 1% of their enclosing space (The MICrONS Consortium 2025). Measuring many inputs to an individual neuron thus requires efficient sampling of tiny synapses distributed within a large volume.

Another challenge is the short time scale of single neuron computation. Neurons integrate inputs over ~10 ms, necessitating sampling rates on the order of 100 Hz (Koch et al. 1996). Conventional two-photon microscopes scan a focus sequentially through every voxel in a volume, typically hundreds of millions for a dendritic arbor. Independent measurements are limited to < 100 MHz by the fluorescence lifetime of indicators (Peron et al. 2015; Yasuda et al. 2006). The resulting volume scanning rate (< 1 Hz) is far too slow to track neural computation.

Random access two-photon microscopy overcomes these challenges by sampling only relevant voxels (Katona et al. 2012; Otsu et al. 2008; Salomé et al. 2006; Iyer et al. 2006). Random access imaging records from arbitrary target voxels at a rate that depends on the number of targets rather than the entire enclosing volume. Random access imaging is faster and more power-efficient than raster scanning when few voxels generate relevant signals. However, acousto-optic deflector (AOD)-based random access imaging suffers from an *access time cost* of ~20–30 µs to jump between targets (Fernández-Alfonso et al. 2014), thousands of times longer than the minimum voxel measurement time (limited by fluorescence lifetime, 2–3 ns).

To record from arbitrary locations in three dimensions at unprecedented speeds, we developed a Random Access Projection Microscope (SLAP2) with a novel scan system having no per-target access time cost. *Projection microscopy* refers to methods that programmably combine multiple pixels into individual measurements (Kazemipour et al. 2019). Compared to Scanned Line Angular Projection microscopy (SLAP) (Kazemipour et al. 2019), SLAP2 combines pixels into measurements more flexibly and does not rely on computational image reconstruction. Using SLAP2, we characterized input-output relationships of individual layer 2/3 (L2/3) pyramidal neurons in mouse primary visual cortex. We demonstrated dual-plane glutamate imaging of hundreds of synapses at ~200 Hz, three-dimensional (3D) recordings of >800 synapses at >50 Hz, and dual-plane membrane voltage recordings of arbitrarily arranged dendritic compartments at 2–10 kHz. These measurements revealed rich relationships between the inputs and output of neurons, including specificity of input tuning, amplification of specific activity features from input to output, and spatial and temporal correlations among synapses encoding these amplified features. SLAP2 is a powerful new tool for studying neuronal computations in the living brain.

## Results

### Time-efficient random-access imaging

SLAP2 combines high-speed line scanning with programmable digital micromirror devices (DMDs) (Hornbeck 1996) to generate spatiotemporal light patterns not achievable with other scanning systems (Figure 1). A laser line focus, extended in the *y*-axis, is swept along the *x*-axis at 10.8 kHz by a polygonal mirror scanner. This rectangular field of view (FOV) illuminates a DMD, which is imaged onto the sample. The 2D DMD array of tilting mirrors, which define the pixels of the image, can each be set to keep (ON) or discard (OFF) light to flexibly shape illumination (Figure 1b). Because the scanned line illuminates the DMD one column at a time, all ON pixels in a given column are simultaneously excited within the sample. As in conventional two-photon microscopes, fluorescence is measured by one detector for each color channel, resulting in one measurement per column per scan. The DMD pattern is updated after each scan. Images are assembled by combining measurements from multiple scan patterns. Because different columns can have different ON pixel patterns, the beam can jump or be reshaped arbitrarily along the *y*-axis as it is scanned, giving SLAP2 random access imaging capabilities without per-target access time cost.

**Figure 1:**
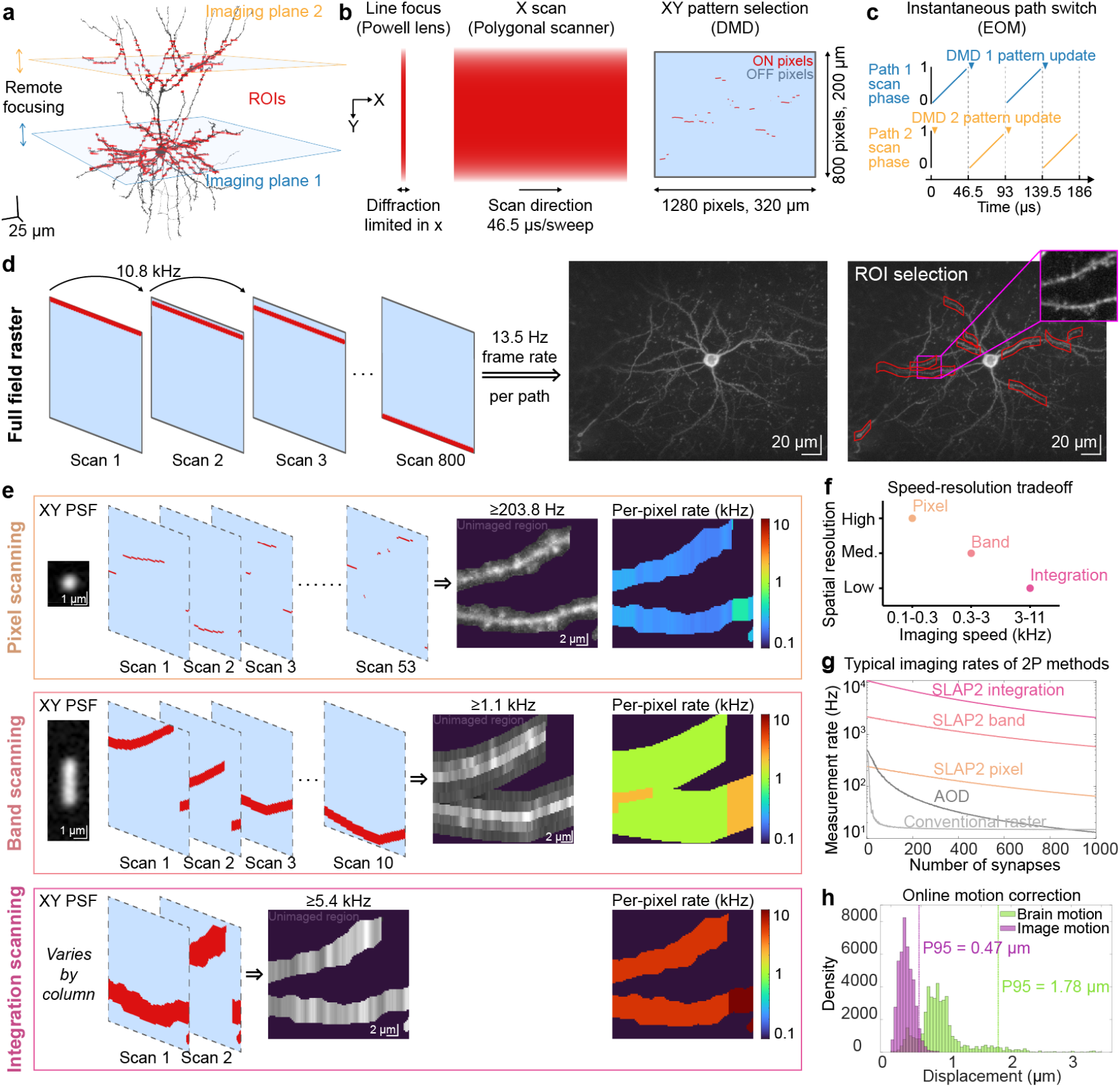
SLAP2 imaging principles. **a**, SLAP2 scanning pattern schematic overlaid on a layer 2/3 pyramidal neuron from The MICrONS Consortium 2025. The two imaging paths sample from two individually-steered planes that scan across depth by remote focusing, and arbitrary ROIs (red) at each depth are imaged using the scanning system described in following panels. **b**, Laser scanning system. A powerful laser is shaped into a high-aspect-ratio line focus by a Powell lens, and switched between two imaging paths. For each path, the illumination line is scanned across a DMD at 10.8 kHz. For each scan, the DMD selects a programmable subset of pixels that are imaged into the sample (ON) and discards remaining illumination light (OFF). **c**, After one path is scanned, the beam is instantaneously switched to the other path with an EOM while the DMD pixel pattern is updated. This results in >99% duty cycle. **d**, Conventional raster scanning is achieved by turning on one row of DMD pixels per scan. (Right) Example raster image (intensity-normalized, gamma = 0.3). Arbitrarily-shaped ROIs (red, overlaid on a full-field raster image) can then be selected for random-access imaging in any combination of three modes: pixel, band, or integration scanning. **e**, Three random access imaging modes demonstrated by zoomed-in DMD patterns and simulated resulting images of the inset in (d) top-right. Excitation PSF in XY and per-pixel measurement rates for each mode are also presented. In pixel scanning, images are recorded at full resolution (with diffraction-limited PSF), and the measurement rate of each pixel is 10.8 kHz/# pixels in column. In Band mode, the Y-dimension of the PSF is elongated, resulting in ROIs composed of uniform-height strips (here, five each), and the measurement rate of each superpixel is 10.8 kHz/# superpixels in column. The height of each strip is fixed, enabling correlation-based motion correction against a reference image acquired with the band PSF. In Integration mode, Y-resolution within each ROI is lost, and the measurement rate of each superpixel is 10.8 kHz/# superpixels in column. The grayscale images are intensity-normalized with gamma = 0.5. **f**, Tradeoff between imaging rate and spatial resolution among SLAP2 scanning modes. **g**, Mean achievable imaging rates of the three SLAP2 modes, AOD-based random access imaging, and resonant raster scanning in simulations involving varying numbers of synapses arranged onto dendritic branches randomly placed within the FOV. **h**, Histogram of total 3D brain displacements (green) and of residual image displacements (magenta) not compensated by online motion correction for a typical acquisition of iGluSnFR4-labeled dendrites at 200 Hz in a mouse running on a wheel.

SLAP2 incorporates two independent imaging paths (Figure S1a). A single polygonal scanner is used for both paths, scanning the two DMDs interleaved, with an electro-optic modulator (EOM) switching between them (Figure 1c). While one DMD is scanned, the pixel pattern on the other is updated, producing a >99% duty cycle. The two paths are independently steered in depth by two rapid remote focus systems (Botcherby et al. 2008), enabling, for example, simultaneous imaging of a neuron’s apical and basal dendrites.

#### SLAP2 imaging modes

SLAP2 performs conventional raster scans by turning ON rows of DMD pixels in sequence (Figure 1d). The intersection of the y-extended excitation line and each x-extended ON pattern results in a point scanned along x. The resulting images have diffraction-limited resolution at numerical aperture≥ 0.82 (2.27± 0.14 µm axial FWHM, mean ± SEM) throughout the 3D FOV (Figure S1b), <17% relative standard variation in brightness across the FOV (Figure S1c), very low excess noise (Figure S1d), and negligible field curvature (<20 nanometers absolute across the imaging field, see **Field curvature**). We next describe three novel random access imaging modes enabled by SLAP2 (Figure 1e), which demonstrate flexible tradeoffs between resolution and speed (Figure 1f).

Pixel scanning: SLAP2 can scan arbitrary regions of interest (ROIs) within the FOV at full spatial resolution. The achievable plane rate in each column is the line rate divided by the number of selected pixels in that column. In contrast, conventional raster scanning microscopes scan the entire bounding box enclosing all regions of interest. In simulated imaging experiments, with varying numbers of synapses organized onto dendritic branches, pixel scanning produced >5-fold speed increases over existing technologies when imaging 400 or more synapses simultaneously (Figure 1g, Figure 2).

**Figure 2:**
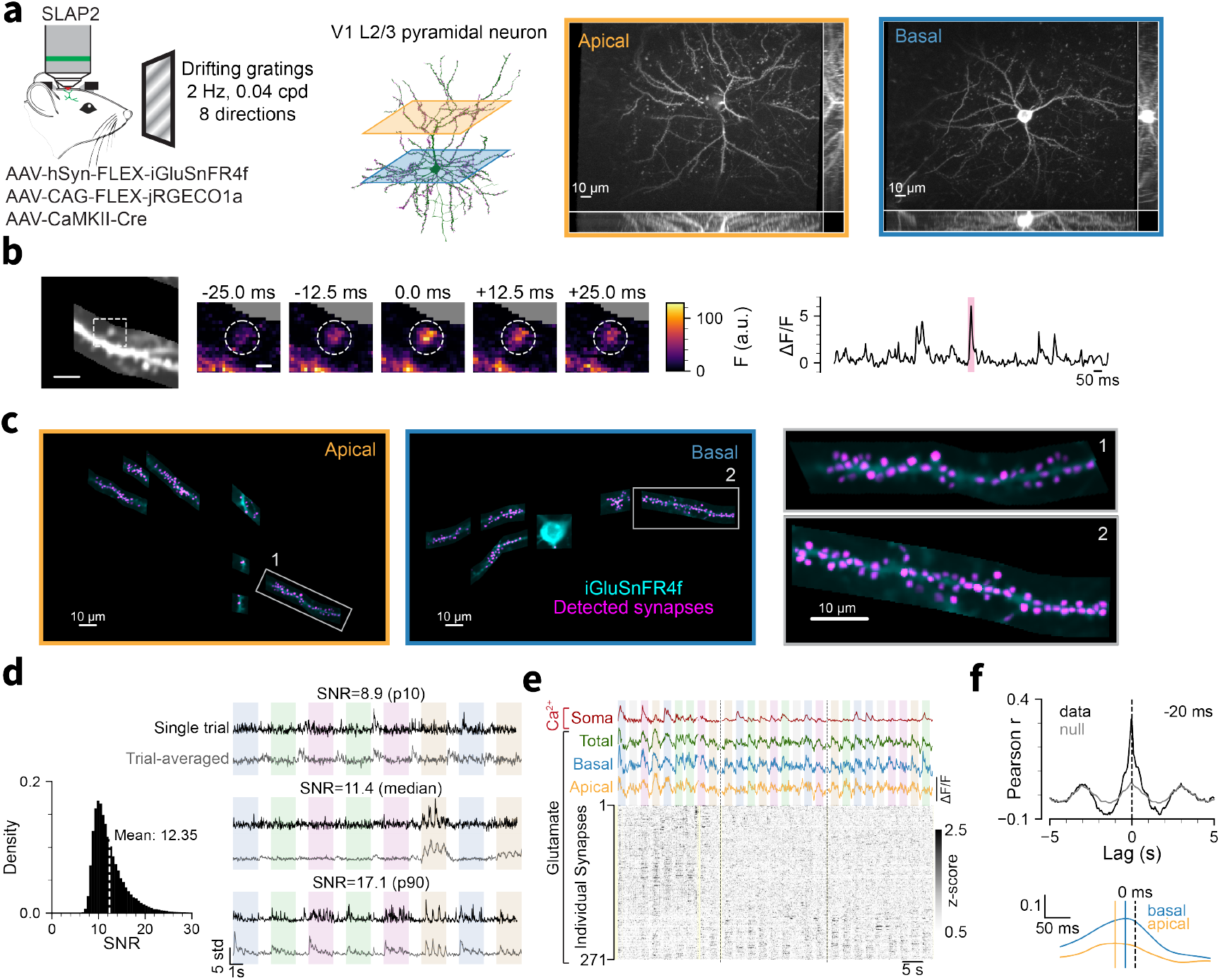
Simultaneous measurements of synaptic inputs and somatic outputs of single neurons with pixel scanning. **a**, Layer 2/3 pyramidal neurons in the primary visual cortex expressed iGluSnFR4f to image synaptic glutamate, and jRGECO1a to image cytoplasmic calcium. Imaging was performed across planes intersecting with apical (orange) and basal (blue) dendrites, while presenting drifting grating stimuli. Images are XY projections of reference stack, with XZ and YZ projections in margins. (scale bar, 10 µm). **b, left:** small dendritic segment (scale bar, 5 µm). **middle:** iGluSnFR4f fluorescence across time, showing fluorescence changes corresponding to glutamatergic transmission at an individual synapse (scale bar, 1 µm). **right:** Fluorescence trace from the synapse; the red highlight indicates the frames shown at left. **c**, pixel scanning. Dendritic segments in the apical (**left**) and basal (**middle**) imaging planes (corresponding to images in **a**) were imaged simultaneously. Synaptic inputs (magenta) were identified based on iGluSnFR4f fluorescence signals (b, d). Magnified view of two dendritic segments with detected synapses (**right**). **d, left:** distribution of signal-to-noise ratio (SNR) for detected synaptic iGluSnFR4f signals across all sessions (*N* = 20). **right:** single-trial and trial-averaged traces from synapses with single-trial SNR at the 10th, 50th, and 90th percentiles of the distribution. **e**, Traces showing somatic calcium (red), glutamate signals summed across apical (orange) or basal (blue) planes, or both (green). The raster (black) shows the activity of 271 individual synapses (sorted using Rastermap (Stringer et al. 2024)) (three consecutive trials, imaged at 200 Hz, 20.5 mW at the sample). **f, top:** For an example neuron, cross-correlation between summed synaptic input and somatic output (black), and the control (gray) computed by shuffling trials within each stimulus direction. The peak at *−*20 ms indicates that output lags input. **bottom:** same analysis separately for apical (orange) and basal (blue) inputs.

Band scanning: To increase plane rates, multiple adjacent DMD pixels within a column can be set ON simultaneously, elongating the PSF in the *y*-axis and reducing the number of scans needed to image a given area. The resulting multi-kHz plane rates enable multiple Z-planes to be scanned at volume rates comparable to single-plane scanning. Using measured point spread functions, we optimized the number, height, and spacing of bands to maximize sampling rates, capture the full width of spiny dendrites, and preserve the ability to distinguish sources separated by one micrometer (Figure S4). The achievable plane rate in each column is the line rate divided by the number of bands in that column, approximately 9 times faster than pixel scanning for the band parameters used here.

Integration scanning: Still higher sampling rates can be achieved by simultaneously illuminating all pixels in each column of a given ROI, thereby integrating pixel columns and discarding all within-ROI vertical resolution. This enables very rapid recordings of the fluctuating brightness of extended objects, especially useful for voltage imaging of neuronal compartments. The achievable imaging rate in each column is the line rate divided by the number of ROIs in that column. Integration scanning retains multi-kHz average imaging rates for over 1000 randomly-distributed small targets across the FOV (Figure 1g, Figure 6).

#### Real-time motion correction

High-resolution random access imaging in the intact brain depends on real-time 3D motion correction (Griffiths et al. 2020). Otherwise, even small sample movements result in lost data or artifacts, as sources move in and out of the sampled regions. SLAP2 uses incoming image data to correct for movement in real time through DMD and remote focus controls, without need for dedicated fiducial objects. A volumetric full-field raster stack is first acquired and used both for ROI selection and as the template for motion correction. During pixel scan imaging of dendrites at ~200 Hz in running mice, residual 3D displacements were <470 nm (95th percentile; Figure 1h). Band scanning produced online corrections with similar performance (Figure S3).

### Input-output transformations measured with pixel scanning

We applied SLAP2 pixel scanning to simultaneously record inputs and outputs of L2/3 pyramidal neurons in the primary visual cortex of awake, head-restrained mice, while presenting visual stimuli to the contralateral eye. We presented eight directions of drifting gratings (2 s, 0.5 s/cycle, 0.04 cycles/degree) in randomized order, repeated at least 40 times each, separated by 1 second of blank screen. Neurons were sparsely transduced with the yellow-green glutamate indicator iGluSnFR4f (Aggarwal et al. 2026) to record synaptic inputs and a red calcium indicator, jRGECO1a (Dana et al. 2016) or RCaMP3 (Yokoyama et al. 2024), to record spiking output signals at the soma. Recordings were made in two planes: one containing the soma and perisomatic dendrites and a second in the apical or lower basal dendrites (Figure 2a).

Release of synaptic glutamate causes brief, localized increases in iGluSnFR fluorescence (iGluSnFR4f decay time 26 ms (Aggarwal et al. 2026)) (Figure 2b–d, Movie S1). On this timescale, synapses whose images overlap in space are rarely coactive, as cortical excitatory neurons have low (<10 Hz) spike rates (O’Connor et al. 2010; Niell & Stryker 2008). Leveraging this spatiotemporal sparsity, we used algorithms inspired by single molecule localization microscopy (Xie et al. 2026; Lelek et al. 2021) to localize putative glutamate release sites with high spatial precision (Figure 2b–c), then used constrained non-negative matrix factorization (Pnevmatikakis et al. 2016) to extract the associated time series (see **Methods**)(Figure 2d, e). These sites, which we refer to as synapses, correspond to synapses identified post hoc in histology (Xie et al. 2026). We detected 0.43 synapses per µm of dendrite. Individual recordings detected up to 398 synapses (median: 195), with mean imaging rates of ~200 Hz and mean SNR of 12.3 (Figure 2d; 20 sessions, 7 mice). The cross-correlation between summed synaptic input and somatic output peaked at −19.4 ±5.6 ms (mean± SEM), consistent with the causal effect of glutamate input on spiking output (Figure 2f) (17 of 20 sessions, 7 mice; 3 outlier sessions excluded, see **Methods**).

#### Selective amplification of stimulus features from input to output

Most recorded somatic outputs (17 of 20 sessions; soma Ca^2+^) and most synaptic inputs (78± 31% per session, mean ±SD) exhibited stimulus-modulated activity (see **Methods**). Synapses exhibited diverse timecourses of peri-stimulus activity, which we refer to as *tuning profiles* (Figure 3a). We used non-negative matrix factorization (NMF) to decompose input tuning profiles into ten factors (nine related to stimulus timing and one baseline offset) per stimulus. The factors captured >94% of the signal energy of the input tuning profiles. For subsequent analyses, we grouped the stimulus timing factors into four temporal categories, corresponding to stimulus onset (*ON*, 2 factors), stimulus offset (*OFF*, 2 factors), grating phase (*CYC*, 4 factors), and post-stimulus ramping associated with suppression by the grating stimuli (*SUP*) (Figure 3b, Figure S5c–d). Individual synapses showed loading onto multiple categories (Figure 3a). Correlations of loadings within and across categories (Figure 3c) were structured. CYC and ON loadings were weakly correlated with each other and anti-correlated with OFF and SUP, whereas OFF and SUP loadings were also weakly correlated with each other. Category loadings were correlated across stimulus identities (*e*.*g*., synapses with OFF responses to one stimulus tended to have OFF responses to other stimuli). Orientation selectivity (Niell & Stryker 2008; Hubel & Wiesel 1962), indicated by strong off-diagonal bands in the correlation matrix, was present for ON and CYC categories, but not for OFF or SUP.

**Figure 3:**
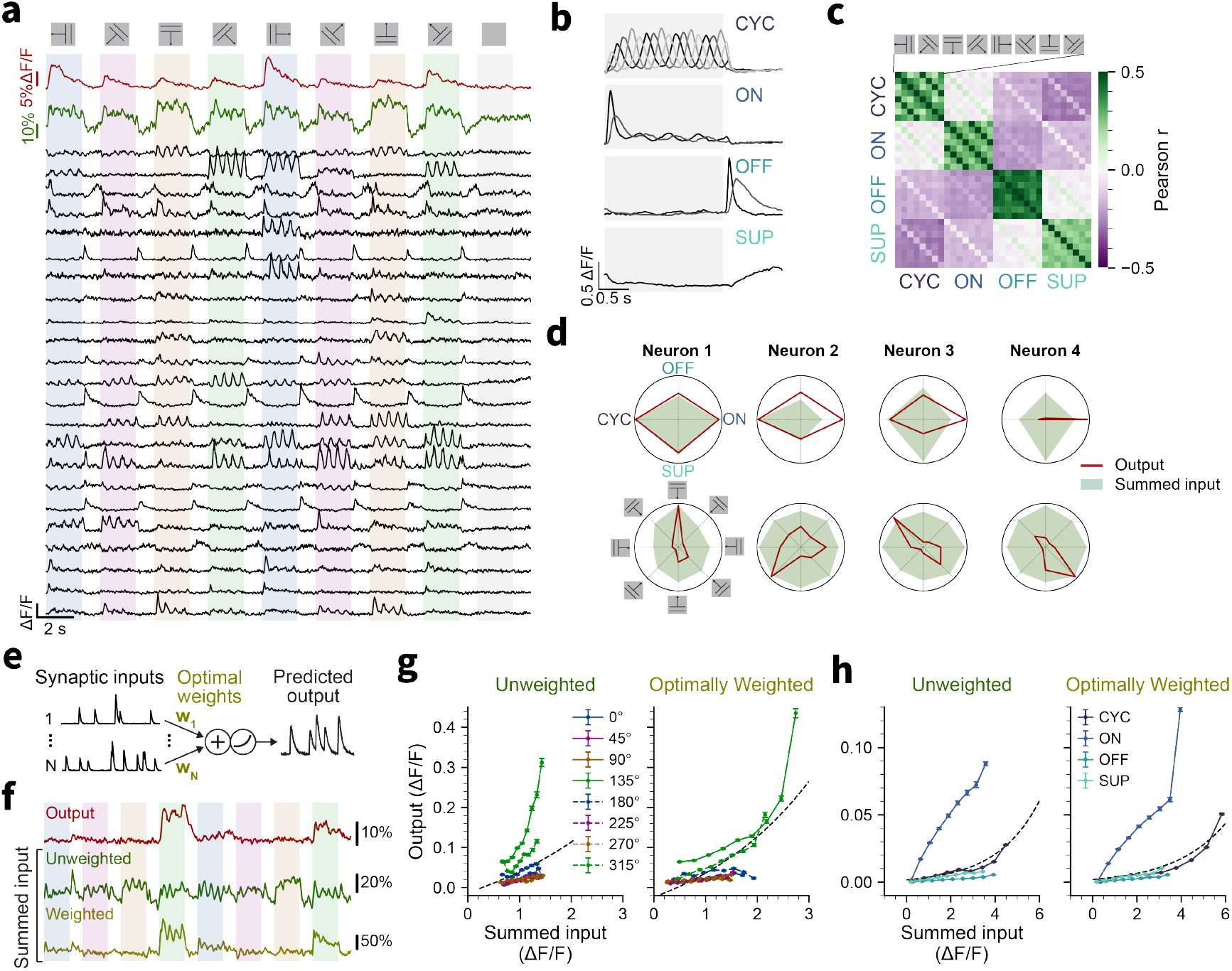
Selective amplification of tuning features from input to output. **a**, Tuning profiles of 23 representative example synapses, summed input (green) and somatic calcium (red). **b**, Non-negative matrix factorization (NMF) of synaptic stimulus responses produced temporal factors which were grouped into four categories: CYC (cyclic responses to grating phase), ON (stimulus onset), OFF (stimulus offset), and SUP (suppression) explaining >94% of the signal energy. **c**, Correlation matrix (Pearson’s *r*) of NMF loadings for each factor category and stimulus identity. **d**, Polar plots showing temporal-factor tuning (top; ON, CYC, OFF, and SUP) and direction tuning (bottom) for summed synaptic input and somatic output in four example neurons. Neurons are sorted by the sum of squared differences between input and output temporal-factor tuning, illustrating selective amplification of specific stimulus directions or temporal response motifs. **e**, Generalized Linear Model (GLM) used to fit a single weight to each synapse. These weights are used to compute optimally weighted summed inputs shown in **f-h. f**, Tuning profiles of somatic output (red), unweighted summed input (green) and optimally weighted summed input (olive) for an example neuron. **g**, Stimulus-dependent activation functions (somatic output vs. summed synaptic input) for unweighted (**left**) and optimally weighted (**right**) summed synaptic input for the neuron in **f**. Dashed line, best fit to the pooled data across all stimulus directions. **h**, Same as **g**, but activation functions are computed separately for each temporal factor category (CYC, ON, OFF, SUP).

Temporal categories were approximately equally represented in the summed input. However, the ON temporal category accounted for a larger proportion of output activity than of input (summed input: 0.20–0.28, median 0.23; output: 0.17–0.83, median 0.32; *p* = 0.003 Wilcoxon signed-rank test) indicating selective amplification of the ON temporal category (Figure 3d, Figure S5e–f). Individually, 10 of 17 stimulus-modulated neurons showed this amplification of the ON proportion (output exceeded input by > 3 SD of the input proportions across sessions; Figure S5f). Separately, the orientation selectivity index (OSI), a measure of orientation tuning, was higher for somatic output than for summed inputs (OSI: soma = 0.47 ±0.05 vs. summed input = 0.15 ±0.03, mean ±SEM, *N* = 17, Wilcoxon signed-rank test, *p* < 0.001), indicating selective amplification of specific input orientations (Priebe & Ferster 2008) (Figure 3d, Figure S5a–b). These results identified two distinct representations, i.e. ON preference and stimulus identity preference, that were selectively amplified from glutamate inputs to somatic output. To investigate these amplifications we characterized neuronal activation functions, which map summed input to the mean corresponding output, computed for different stimulus identities (Figure 3g) or temporal categories (Figure 3h). For a given feature, mean output increased monotonically with input. However, outputs were higher for a given input during preferred vs non-preferred stimuli and for the ON factor vs other factors, indicating that a given amount of presynaptic glutamate release is more effective at driving output for the amplified temporal features (Figure 3g-h).

Amplification could arise from synapses tuned to these features having larger postsynaptic weights. To evaluate this possibility, we fit generalized linear models (GLMs), which sum weighted observed synaptic inputs and a bias then apply a single nonlinearity to optimally predict output. The resulting weighted inputs accounted for only a small fraction of the observed amplification (Figure 3e-h, Figure S5j-k). In contrast, for data-driven simulations in which amplification arose from postsynaptic weights proportional to each synapse’s preference for the amplified feature, GLMs accounted for amplification, even when only 5% of synapses were observed (Figure S5g-i). These analyses suggest that amplification does not arise from a single nonlinearity or the simple weighting of synapses proportional to their tuning preference, and that additional factors contribute to the transformation of tuning features from input to output.

#### Spatiotemporal organization of synaptic inputs

Input-output amplification could also arise from spatiotemporal organization of inputs (Spruston 2008; London & Häusser 2005; Kerlin et al. 2019). For example, currents originating far from the soma may undergo dendritic attenuation and have a smaller effect on somatic potentials compared to proximal inputs. Synapses that form functional clusters could integrate nonlinearly by recruiting active dendritic conductances (Major et al. 2013).

In ten sessions we collected large structural stacks of the neuron and manually reconstructed the dendritic arbor, allowing us to measure distances (cable and Euclidean) between synapse pairs and between synapses and the soma (Figure 4a, Figure S6d).

**Figure 4:**
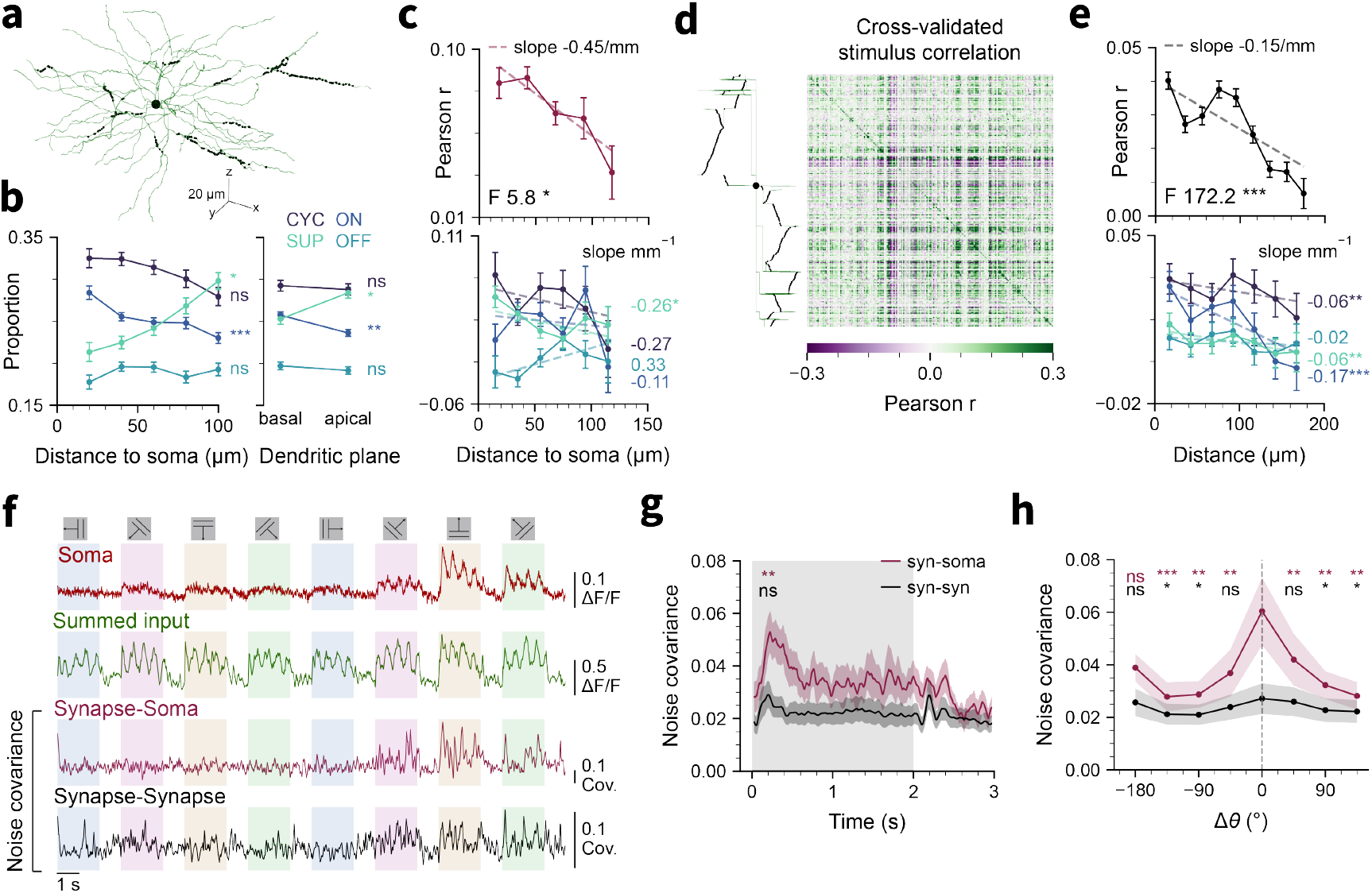
Spatiotemporal organization of synaptic inputs and somatic outputs. **a**, Reconstruction of a V1 L2/3 pyramidal neuron with imaged synapse locations mapped onto the dendritic tree. **b**, Proportion of each temporal factor category (ON, CYC, OFF, SUP) per synapse, as a function of Euclidean distance to the soma (**left**; *N* = 10 sessions, ANCOVA) and location on basal vs. apical dendrites (**right**; *N* = 8 sessions with both apical and basal synapses imaged, two-way ANOVA). **c**, Cross-validated stimulus correlation between synapses and the soma as a function of Euclidean distance to the soma (**top**) and separately for each temporal factor category (**bottom**) (ANCOVA, *N* = 10 sessions). **d**, Pairwise cross-validated stimulus correlations between synapses (**right**), ordered by anatomical location on the dendritic tree (dendrogram, **left**). **e**, Cross-validated stimulus correlation between synaptic pairs as a function of Euclidean distance (**top**), and separately for each temporal factor category (**bottom**) (ANCOVA, *N* = 10 sessions). **f**, For an example neuron, tuning curves of somatic calcium (red), summed input (green), and noise covariance between soma and synapses (purple), and noise covariance between synaptic pairs (black) averaged across synapses. **g**, Synapse-soma (purple) and synapse-synapse (black) noise covariance as a function of time relative to stimulus onset (mean*±* SEM, *N* = 17 sessions; Wilcoxon signed-rank test comparing mean during 0.0–0.5 s and 0.5–2.0 s windows). **h**, Synapse-soma (purple) and synapse-synapse (black) noise covariance as a function of difference in preferred direction relative to the soma (Δθ) (paired Wilcoxon signed-rank test against value at Δθ = 0; *N* = 15 sessions with soma OSI > 0.3). (* *p* < 0.05, ** *p* < 0.01, *** *p* < 0.001)

We first investigated whether synapse tuning is spatially organized relative to the soma. The proportion of ON input was larger for synapses closer to the soma (ANCOVA, *F* = 11.4, *p* < 0.001), whereas the proportion of SUP was larger farther from the soma (ANCOVA, *F* = 5.9, *p* = 0.015) (Figure 4b). Similarly, the proportion of ON input was larger in perisomatic dendrites (two-way ANOVA, *F* = 7.1, *p* = 0.008), and the proportion of SUP was larger in apical dendrites (two-way ANOVA, *F* = 4.1, *p* = 0.04). We quantified tuning profile similarity by computing *cross-validated stimulus correlation* (CVSC).

Tuning profiles used to compute CVSCs were drawn from non-overlapping sets of trials, eliminating contribution from shared noise (Cohen & Kohn 2011). Synapses closer to the soma had higher CVSC with the soma (slope −0.45 mm ^*−*1^; ANCOVA, *F* = 5.8, *p* = 0.02) (Figure 4c). We did not detect any significant spatial organization of noise correlations (correlations in residuals after subtracting mean stimulus response) (ANCOVA, *F* < 1, *p* = 0.9) (Figure S6a).

We next investigated whether functional properties of synapses are spatially clustered (Wilson et al. 2016). CVSCs between synapse pairs decreased with distance (ANCOVA, *F* = 172, *p* < 0.001, slope −0.15 mm ^*−*1^; Figure 4d–e). Computed for stimulus preferences within temporal categories, this effect was stronger for ON than other factors (ANCOVA, *F* = 86, *p* < 0.001, slope− 0.17 mm ^*−*1^) (Figure 4e). Noise correlations between synapse pairs showed a similar distance-dependent decrease (ANCOVA, *F* = 5548, *p* < 0.001) (Figure S6a). Synapse-soma and synapse-synapse clustering were both larger and more statistically significant when using Euclidean rather than cable distance, although results using cable distance were similar (Figure S6a).

Neuronal activation is influenced not only by the mean, but also by the variance of inputs (Salinas & Sejnowski 2000). We therefore evaluated whether covariance of inputs is stimulus-modulated and its relationship to output. We computed cross-trial noise covariance among synapses and between synapses and the soma (Figure 4f). Synapse-soma covariance was significantly higher during the stimulus onset window (0–0.5 s, corresponding to the ON temporal factor category) than during the remainder of the stimulus (0.5 s–2 s) or the post-stimulus period (2–2.5 s) (Figure 4g, *p* < 0.01). Synapsesoma covariance was also significantly higher for the neuron’s preferred direction compared to other directions. Synapse-synapse covariance showed weaker dependence on peri-stimulus timing or direction, but was significantly larger for the preferred direction than nonpreferred ones (Figure 4g,h). These results show that features that were amplified by neuronal input-output transformations exhibited specific spatial and temporal organizations of synaptic inputs. ON-preferring synapses that shared the neuron’s stimulus preferences were preferentially located near the soma, and were spatially clustered. Trial-to-trial fluctuations in input were more strongly correlated to corresponding fluctuations in somatic output for the amplified stimulus features. Together, these results demonstrate capabilities of SLAP2 for investigating neuronal computations and the precise temporal and spatial organization of synaptic input activity.

#### Predicting somatic output with GLMs

Theoretical work has suggested that linear integration with a single nonlinearity, modeled by GLMs, can explain much of a simulated L2/3 neuron’s voltage response (Ujfalussy et al. 2018), but data to constrain models under naturalistic conditions has previously not been available.

We built GLMs that predict somatic calcium output from the stimulus timing (which can account for information from unobserved inputs) and varying numbers of the observed synaptic inputs (Figure S7a– b). Model performance on held-out test data improved as increasing numbers of synapses were included as predictors (Figure S7c). A model using only the summed glutamate input and stimulus context as predictors performed worse than models with individually-weighted synapses (Figure S7d). A GLM constrained to use positive, temporally decaying weights for each synapse (reflecting properties of excitatory synapses) predicted output as well as the unconstrained GLM (Figure S7f). The fit weights for both GLMs were larger for synapses with similar tuning to the soma, despite inclusion of the unregularized stimulus timing predictor that captures mean tuning effects, indicating that trial-to-trial variations of these synapses better predicted variations in somatic output than other synapses (Figure S7e, g).

### Broader sampling of synapses with volumetric band scanning

Given the capability of SLAP2 to sample large numbers of targets, we next sought to understand whether input-output predictions saturate as more synapses are recorded. To push the sampling abilities of SLAP2, we used volumetric band scanning to sample more targets along a neuron’s dendritic tree. Band scanning achieves plane rates of up to 2.2 kHz per path, enabling volumetric imaging by using the remote focus systems to scan in depth.

We evaluated band scan signal quality by interleaving pixel and band scan recordings of the same iGluSnFR4s-expressing dendrites (imaging rate, ~200 Hz) (Figure S8a,c). In these recordings, ~90% of time was devoted to pixel scanning, and ~10% to band scanning. Despite the lower *y*-resolution of band scanning, sources of glutamate activity visible during pixel scanning could also be detected from the band scanning data (Figure S8d–g). Across four datasets from two mice, 74.3% of the sources identified by pixel scanning could be matched to a source detected by band scanning, with high SNR (band scan mean, 8.27; pixel scan mean, 11.97) (Figure S8h). OSI was calculated for each source, and for pixel scan sources that were orientation-selective (OSI>0.5), their preferred orientations differed from that of their matched band scan sources by <23.3° (90th percentile), approximately half the interval between grating orientations (Figure S8i–j).

We performed volumetric band scanning on neurons expressing iGluSnFR4s and RCaMP3 in V1 with drifting grating stimuli, at volume rates sufficient to capture iGluSnFR4s transients (53–121 Hz, iGluSnFR4s decay time 153 ms (Aggarwal et al. 2026)) (Figure 5a–e, Movie S2). In recordings from 6 distinct neurons from 3 mice, we identified up to 970 (median: 834) glutamate sources per neuron (mean SNR, 8.41) (Figure 5f–g). An average of 0.68 sources per micron of dendrite were detected. Extracted sources showed diverse tuning preferences similar to our pixel scan recordings (Figure 5h). We built GLMs to predict somatic output from recorded glutamate inputs in band scanning data (see **Methods**). As with pixel scanning data, a GLM using the total glutamate signal performed worse than the model with all synapses as individually-weighted inputs (Figure 5i). We also found that performance continued to improve with synapse counts beyond those observed in pixel scans (Figure 5j), indicating that the inputs that predict neuronal output in V1 are high-dimensional, even when only low-dimensional visual stimuli are presented.

**Figure 5:**
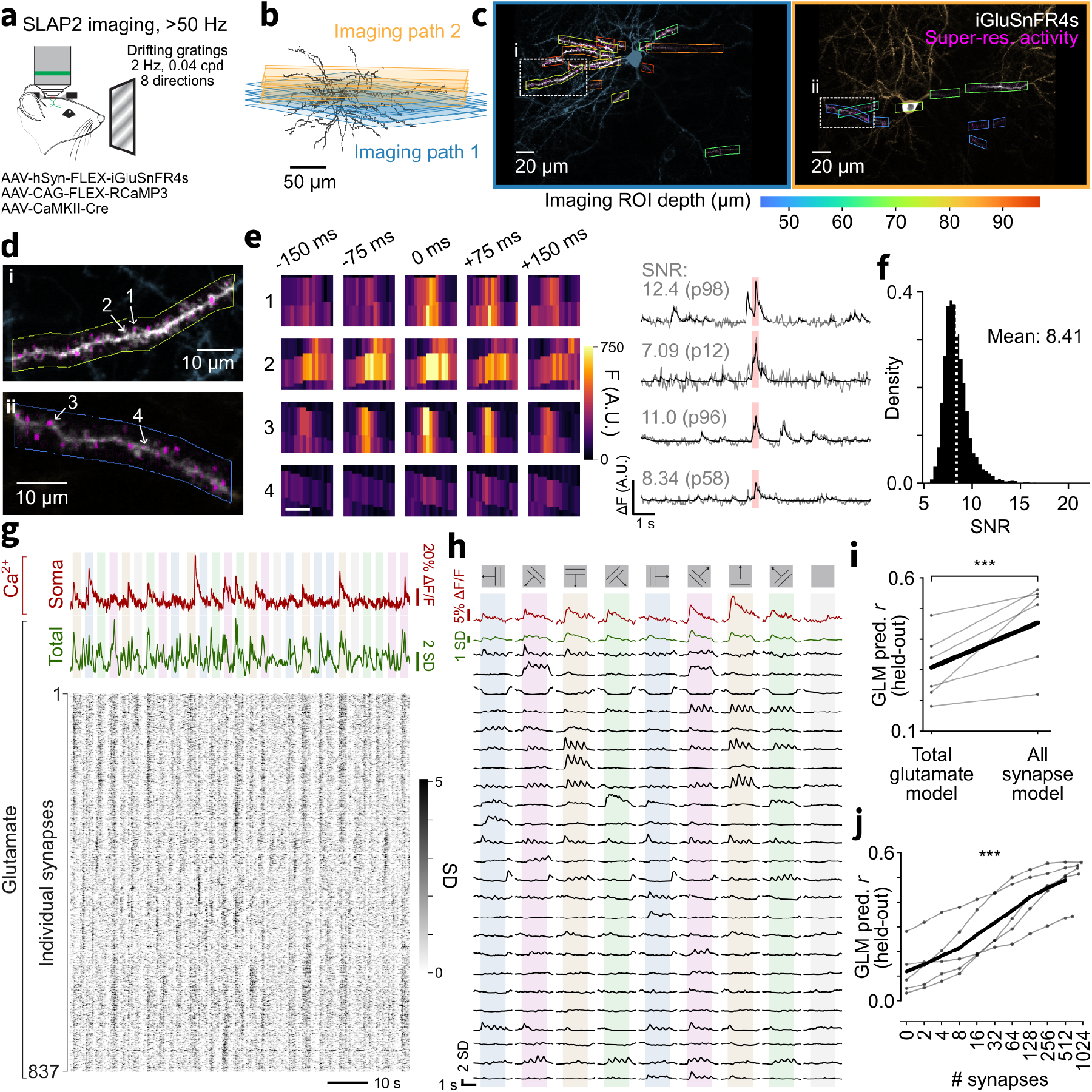
Volumetric synaptic imaging using band scanning. **a**, Layer 2/3 pyramidal neurons in the primary visual cortex expressed iGluSnFR4s to image synaptic glutamate, and RCaMP3 to image cytoplasmic calcium. SLAP2 imaging was performed while presenting drifting grating stimuli. **b**, Volumetric imaging was performed by scanning over several planes on each imaging path (blue and orange). **c**, Maximum intensity projection across depth of the reference stack of an imaged neuron. SLAP2 imaging ROIs for band scanning are colored by depth from the pia. The super-resolution activity image is overlaid in magenta. **d**, View of the two dendritic segments and overlaid activity image (magenta) denoted by the dotted boxes in (c). **e, left:** iGluSnFR4s fluorescence across time, showing band scanning measurements corresponding to glutamate transmission at the four synapses marked in (d). Scale bar, 2 µm. **right:** Fluorescence trace from the synapse (least-squares trace in grey and denoised trace in black). Red highlight indicates the time window shown on the left. Signal-to-noise ratio (SNR) calculated on each least-squares trace and the percentile across all sources is listed. **f**, Distribution of SNR of least-squares traces from band scan measured iGluSnFR4s signals across six sessions. **g**, Traces showing somatic calcium (red), glutamate signals summed across all detected sources (green) in three consecutive drifting grating trials (shaded bars denote when the gratings were presented). The raster (black) shows the corresponding activity of 837 individual synapses (sorted using Rastermap (Stringer et al. 2024)). Glutamate traces are plotted in units of noise standard deviation (SD). **h**, Tuning profiles of somatic calcium (red), summed glutamate input (green), and 24 representative example synapses (black). **i**, Comparison of GLM performance between a model that assigns equal weight to all synapses (“total glutamate”) and a model that assigns a separate weight to each synapse (“all synapse”). Mean (black) across six datasets (each in grey) is shown. (*** *p* < 0.001 on paired two-sided t-test) **j**, Generalized linear model (GLM) performance (Pearson *r* of predictions with measurements on held-out data) as a function of the number of synapses included in the model. Mean (black) across the five datasets (each in grey) that had at least 400 total measured synapses is shown. (*** *p* < 0.001 on ANCOVA)

**Figure 6:**
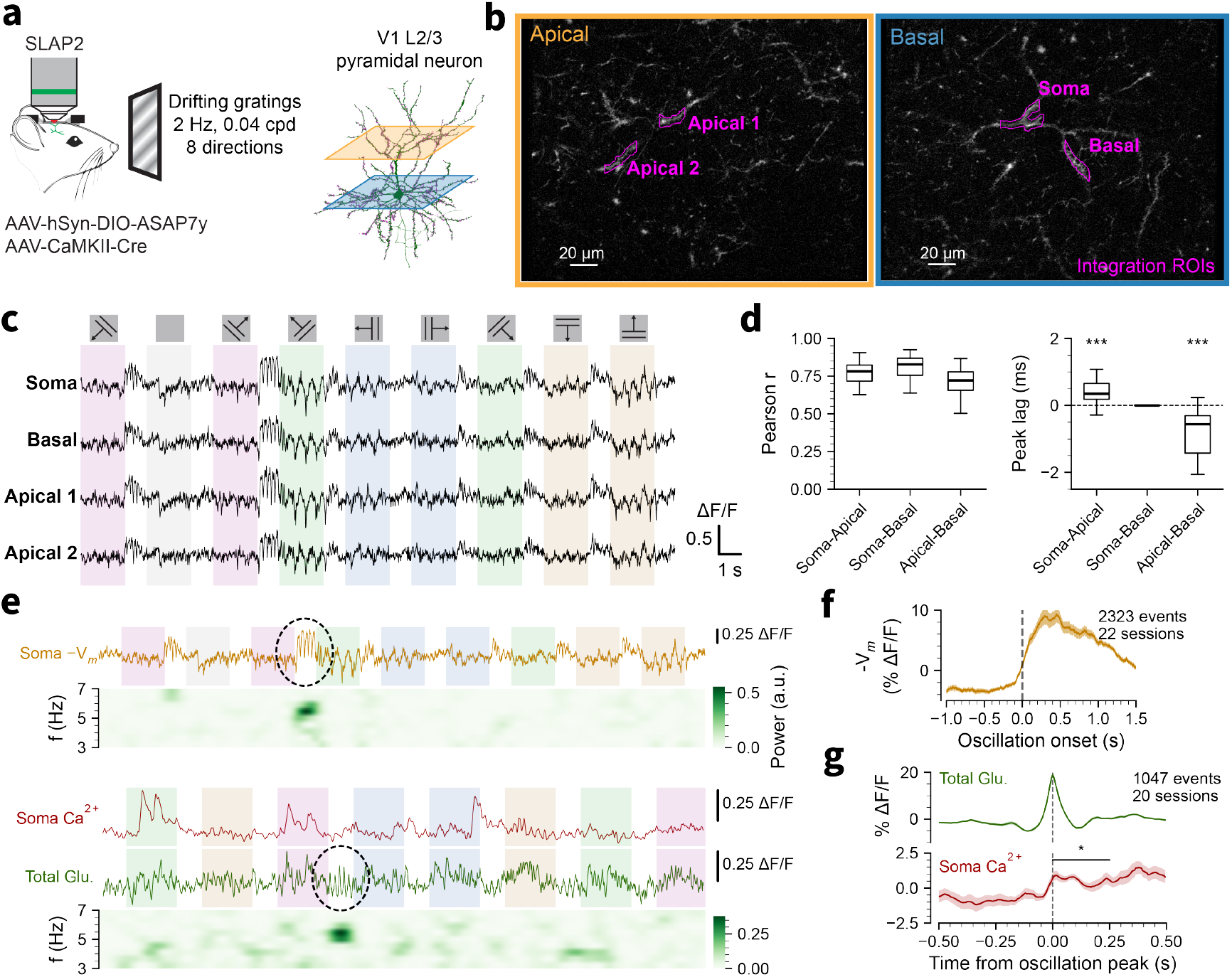
Imaging membrane potential across multiple dendritic compartments. **a**, Layer 2/3 pyramidal neurons in the primary visual cortex expressed ASAP7y. Imaging dendritic membrane potential was performed across multiple apical and basal compartments while presenting drifting grating stimuli. **b**, Integration scanning mode. Dendritic segments (magenta) in the apical (left) and basal (right) imaging planes were imaged. **c**, Traces showing membrane potential activity in somatic, basal and apical compartments corresponding to ROIs shown in **b. d, left:** Correlation (Pearson *r*) between compartment pairs (soma-apical, soma-basal, and apical-basal). **right:** Lag between compartment pairs, computed as the peak lag of pairwise cross-correlations; a positive soma-apical lag indicates that the soma leads the apical dendrite, and a negative apical-basal lag indicates that the basal dendrite leads the apical dendrite (*N* = 22 sessions; one-sample Wilcoxon signed-rank test against zero). **e**, low frequency oscillations (dashed circle) in membrane potential (**top**) and in total glutamate and somatic calcium (**bottom**) traces, with corresponding spectrograms. **f**, Mean membrane voltage aligned to oscillation onset (22 sessions; mean *±*SEM across sessions). Positive Δ*F*/*F* indicates hyperpolarization. **g**, Total glutamate and somatic calcium during oscillatory events, aligned to the oscillation peak (1047 events with a valid somatic trace in trials < 41 across 20 sessions; mean *±* SEM across sessions; paired Wilcoxon signed-rank test over sessions, 0 to 0.25 s vs −0.5 to −0.25 s baseline).

### Imaging of membrane potential across multiple dendrites with integration scanning

The relative amplitude and delay of membrane potentials across a cell’s dendrites can reveal where input synapses reside and the biophysical mechanisms they recruit (Lee et al. 2026; Park et al. 2025; Petersen 2017; Smith et al. 2013). Genetically encoded voltage indicators enable measurements of membrane potential fluctuations with high spatial and temporal resolution (Hao et al. 2026), but stably recording these indicators from many cellular compartments at once at high speed has been a challenge for conventional microscopy.

To evaluate the utility of SLAP2 integration scanning for such measurements, we imaged L2/3 V1 pyramidal neurons expressing the genetically encoded voltage indicator ASAP7y (Hao et al. 2026). We simultaneously recorded from many dendritic compartments across two planes at 2 kHz each (Figure 6a-b; Figure S9a). We observed action potentials and subthreshold modulations tuned to stimuli (Figure 6c; Figure S9b), even from short (<5 µm) dendritic segments across all depths sampled, using <70 µW/µm^2^ of illumination intensity, targeted exclusively to desired structures, at >200 µm below the pia.

Voltage signals across compartments within a neuron were highly correlated, largely independent of whether compartment pairs were on the same or different imaging planes (soma–apical: 0.75 ±0.03, soma–basal: 0.81 ±0.02, apical–basal: 0.71 ±0.02; mean± SEM, *N* = 22 sessions across 5 mice), consistent with the electrotonic compactness of L2/3 pyramidal neurons (Wong-Campos et al. 2023) (Figure 6d). However, soma and basal dendrites led apical dendrites in temporal cross-correlations by 0.5 ±0.14 ms (soma–apical, mean ±SEM; *p* < 0.001, Wilcoxon signed-rank test against zero, *N* = 22 sessions) (Figure 6d), reflecting propagation of voltage signals.

We observed temporally discrete oscillations in these recordings that could be detected as localized peaks in time-frequency spectrograms of somatic and dendritic membrane potential. These events spanned frequencies of 2.6–6.9 Hz (95% interval) and had durations of 1.3 ±0.6 s (mean ±SD; median 1.05 s; *N* = 2323 events across 22 sessions), had strongly hyperpolarized troughs, and preferentially (but not universally) started at the offset of passive visual stimuli (Figure 6e-f, top). Such oscillations have previously been observed using intra- and extra-cellular electrophysiology (Einstein et al. 2017; Senzai et al. 2019; Nestvogel & McCormick 2022). We detected similar oscillations in summed glutamate input in our synaptic recordings (Figure 6e, bottom), with frequencies of 2.5–6.6 Hz (95% interval) and durations of 1.3 ±0.5 s (mean ±SD; median 1.10 s; *N* = 1369 events across 20 sessions). Almost all individual synapses participated in oscillations (Figure S9c-d).

Although oscillations in total input involved many simultaneously-active synapses, somatic calcium showed only weak modulation aligned to oscillation peaks (1.4± 0.6% Δ*F*/*F*, mean± SEM across 20 sessions, in the 0–0.25 s window relative to the −0.5 to −0.25 s baseline; *p* = 0.021, paired Wilcoxon signed-rank test; Figure 6g). This is likely due to strong concomitant inhibitory input revealed by the hyperpolarization observed in our voltage recordings (Figure 6f).

Together, these results demonstrate SLAP2’s ability to perform sensitive subcellular voltage imaging and its potential for multimodal experiments in neuronal computation.

## Discussion

SLAP2 is a random access projection microscope designed to observe the activity of many synapses and neuronal output of individual neurons *in vivo*. SLAP2 derives unique advantages from its flexible scan system, which has no per-target access time cost and can rapidly change the shape of excitation light. Its imaging modes (Figure 1e) trade spatial resolution for speed, while its online 3D motion correction tracks the sample for random access imaging of small structures (Griffiths et al. 2020).

SLAP2 is optimized for more time- and power-efficient imaging of many synapses in behaving animals. AOD-based random access microscopes can rapidly record from small numbers of targets, especially in 3D (Akemann et al. 2022; Nadella et al. 2016; Katona et al. 2012), but their per-target access time limits imaging rates when many targets are sampled (Figure 1g). Bessel-droplet imaging rapidly records projections of dense 3D volumes, but has lower power efficiency and lower frame rates than random access methods (W. Chen et al. 2024).

Limitations of SLAP2 are its fixed 320× 200 µm lateral FOV per imaging path and fixed 1030 nm excitation wavelength. The FOV constraint prevents applications at scales much larger or smaller than an individual neuron’s dendrites, and the scan system is most efficient when structures of interest exist across the entire x-axis. SLAP2 requires a powerful laser (150 W, 50 MHz, 100 fs, 1030 nm) to generate the long illumination line at the Powell lens, and up to >99% of power is discarded at the DMDs, depending on the scanning mode (see **Excitation path**). Suitable commercial lasers only exist at >1 µm wavelengths. The 1030 nm wavelength is nevertheless ideal for single-excitation, dual-emission two-photon imaging of YFP- and RFP-based indicators (Mohr et al. 2020; Aggarwal et al. 2026; Yokoyama et al. 2024; Dana et al. 2016), used here for separate input and output channels. Because most of the input power is discarded at the DMDs, the total laser power delivered to the brain remains below 100 mW regardless of imaging mode, below the threshold for heating-induced damage (Podgorski & Ranganathan 2016).

Advances in commercialized femtosecond lasers at other wavelengths, and DMDs with improved power handling, array size, and switching rates, would enable further applications and improvements. We also envision several extensions of the existing system. First, beyond imaging applications, SLAP2 can illuminate any pixel pattern with only a single line scan, making it highly suitable for high-bandwidth optogenetic stimulation, glutamate uncaging, and two-photon lithography. Second, SLAP2’s highly flexible scan system and real-time data pipeline make it well suited to closed-loop experiments, such as adaptive structural imaging that concentrates light dose on the most informative regions of a sample (Ye et al. 2025). Similarly, ongoing neural activity could determine subsequent imaging or photostimulation patterns. Third, SLAP2’s flexible measurements could enable compressive sensing methods by designing mixed measurements based on image content or sample statistics (Kazemipour et al. 2019; Whang et al. 2025; Li et al. 2024).

Band scanning samples sparse bands with a point spread function that is elongated in *y* to increase imaging rates. To extract synaptic signals from these anisotropic-resolution data, we developed new algorithms that build on prior localization and demixing techniques (Benisty et al. 2022; Xie et al. 2026; T.-W. Chen et al. 2025; Pnevmatikakis et al. 2016; Lelek et al. 2021). We validated these algorithms against interleaved pixel scanning of the same dendrites, a within-recording comparison that illustrates the flexibility of SLAP2’s scan system. Band scanning recovered most sources detected by pixel scanning, along with their stimulus preferences, using a fraction of the measurement time albeit at lower SNR (Figure S8). SLAP2 therefore allows users to trade resolution for coverage according to their experimental question. More generally, SLAP2’s flexible scan system allows other scanning modes to be developed and tailored to specific imaging problems with software alone.

We applied SLAP2 to image L2/3 pyramidal neurons in the visual cortex driven by drifting grating visual stimuli. Consistent with previous dendritic calcium imaging studies in visual cortex (Jia et al. 2010; T.-W. Chen et al. 2013; Wilson et al. 2016; Iacaruso et al. 2017), we found that the synaptic inputs to a single neuron have diverse stimulus preferences, many of which differ markedly from the neuron’s output. Unlike dendritic calcium signals, which are dominated by the postsynaptic neuron’s own activity and from which synaptic components must be isolated (Kerlin et al. 2019; Y. Chen et al. 2026), glutamate imaging reports presynaptic release directly (Kazemipour et al. 2019; Aggarwal et al. 2026; Y. Chen et al. 2026). By recording glutamate from many synapses simultaneously at high speed, we observed rich spatiotemporal structure in inputs and millisecond-timescale lags between inputs and output. Volumetric imaging accessed more dendritic length and recorded from more synapses, improving moment-to-moment prediction of firing output by GLMs, a model class that can explain much of the voltage response of biophysically simulated L2/3 neurons (Ujfalussy et al. 2018). We found evidence for multiple mechanisms shaping computations in these cells. For example, the nonlinearly-amplified ON response was disproportionately represented by synapses nearest the soma, exhibited functional clustering, exhibited larger correlations of stimulus preferences between inputs and output, and was more strongly represented among synapses with larger GLM weights. Notably, ON marks the only moment in our stimulus paradigm where the visual stimulus changes unpredictably.

Integration scanning enabled simultaneous voltage recordings from somatic, basal, and apical compartments across two planes at 2 kHz. Voltage signals across compartments were highly correlated, consistent with the electrotonic compactness of L2/3 pyramidal neurons (Wong-Campos et al. 2023), yet soma and basal dendrites led apical dendrites by ~ 0.5 ms. We also observed strongly hyperpolarizing oscillations in somatic and dendritic membrane potential that preferentially started at the offset of visual stimuli, and similar oscillations in glutamate input recorded in separate experiments, in which nearly all synapses participated. Low-frequency membrane potential oscillations reduce visual responses of L2/3 neurons (Einstein et al. 2017), and a 3–5 Hz rhythm in V1 associated with low arousal, proposed to follow the withdrawal of sustained input, depends on thalamocortical interactions (Nestvogel & McCormick 2022). Because such oscillations also occur across arousal states, with stimulus timing that differs across behavioral conditions (Einstein et al. 2017), whether these phenomena are identical, and how they relate to sensory processing, remains unclear. Combining SLAP2 measurements of membrane potential across compartments with the tuning and trial-to-trial participation of many individual inputs could reveal how these oscillations arise and how they shape sensory processing in pyramidal neurons.

Studies have modeled how individual neurons can implement input-output operations, including hierarchical calculations (Kim et al. 2023; Beniaguev et al. 2021; Jones & Kording 2021; Ujfalussy et al. 2018). Testing such models will require systematic sampling of input space by presenting many distinct stimuli and/or precisely manipulating inputs, for example by glutamate uncaging or closed-loop photostimulation, which were not performed here but are compatible with our methods. Fitting rich models from partial observations of synaptic inputs will likely require new modeling methods and priors from biophysical, anatomical, or functional studies.

In conclusion, SLAP2 addresses critical challenges for high-resolution *in vivo* imaging by enabling precise, programmable control of laser scanning illumination patterns without per-target access time costs, imaging modes defined in software, online motion correction, and an optical design optimized for dendritic imaging. The resulting high-bandwidth input-output measurements enable a new and general approach for studying neuronal computations.

## Data and code availability

Experimental data used for this paper are listed on CodeOcean (https://codeocean.allenneuraldynamics.org/collections/86d03760-b434-4987-ba9e-9489ce47061a), and data can be accessed on S3 at the `aind-open-data` bucket. Motion correction and source extraction for pixel scan glutamate imaging data and motion correction for band scan glutamate imaging data were performed with GIAnT-MATLAB (publicly available on GitHub). Source extraction for band scan glutamate imaging data was performed with GIAnT-Python (publicly available on GitHub). Analysis code for pixel scan and integration scan experiments is deposited at https://github.com/AllenNeuralDynamics/slap2_paper_figures. Other code used for this paper will be made publicly available and listed on CodeOcean prior to publication.

## Acknowledgments

We thank the Lab Animal Services, Neurosurgery & Behavior, Scientific Instrumentation & Process Engineering teams at the Allen Institute for technical support, B. Wynalda and S. Narayan for assistance in cloning, X. Jiang for optical engineering support, and B. Cruz for assistance developing Bonsai workflows.

This research was supported by the Allen Institute, founded by Jody Allen — chair and co-founder of Allen Family Philanthropies, and the late Paul G. Allen — investor, philanthropist, and co-founder of Microsoft. We gratefully acknowledge their vision and generosity, which make this work possible. Work at Janelia Research Campus was funded by the Howard Hughes Medical Institute (HHMI). This research was also supported by the National Institutes of Health (NIH) (DP2NS136990 [K.P.], UM1MH136462 [K.P., K.S., M.Z.L.], R44MH129023 [J.G.], R01EB037653 [A.S.C.], R00MH121533 [M.D.G.], R01NS123681 [M.Z.L.], R01NS131389 [C.C., G.T.]), the United States Department of Defense (PR230899 [C.C.]), Tiny Blue Dot Inc. (MSN196438/AAC1335 [G.T.]), and the Chan Zuckerberg Initiative (CP2-1-0000000704 [K.P., A.S.C.]). The content is solely the responsibility of the authors and does not necessarily represent the official views of the NIH. M.E.X. was supported by the Paul & Daisy Soros Fellowships for New Americans and NIH fellowships F30MH138009 and T32GM136577.

## Author contribution matrix

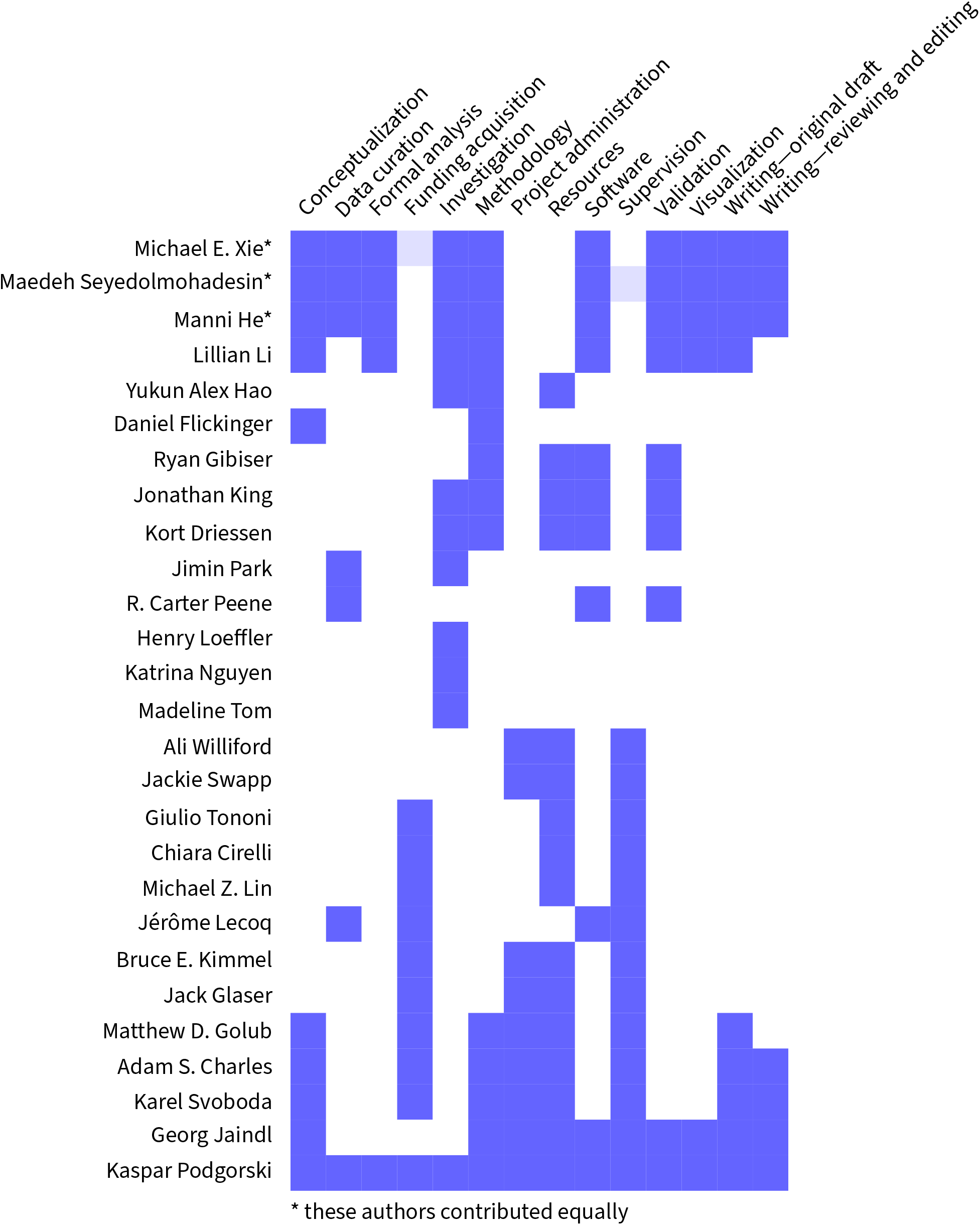

## Methods

### SLAP2 microscope

#### Excitation path

The SLAP2 microscope uses a single powerful laser source (Yb:YAG, 1030 nm, 150 W, 100 fs, 50 MHz; Monaco, Coherent). This high input power is required because the vast majority (>99% during high-resolution scans) of light is discarded by DMD OFF pixels; average power at the sample is typically <70 mW total for raster scanning, or 1–4 mW/ROI for integration scanning at 2 kHz. Pixel scanning recordings were performed at a power density of 6.9 µW/µm^2^, corresponding to total powers of 20– 70 mW, depending on sampling area. The laser includes a module that uses self-phase modulation (SPM) to broaden its spectrum and produce <100 fs recompressed pulses. A dichroic shortpass mirror (14-038, Edmund Optics) rotated away from 45° angle of incidence rejects red-shifted SPM output, reducing the center wavelength of the beam to ~1028 nm and the beam power to ~100 W. A motorized zero-order half-wave plate, high-power laser line polarizing beamsplitter, and water-cooled beam dump (all Thorlabs) control the excitation beam.

A custom-coated high-precision Powell lens (MBF Bioscience) generates the vertically-oriented high-aspect-ratio line beam shared by the two imaging paths. A beam stabilization system stabilizes the beam position by sensing the deviations of a pick-off beam conjugated to the image plane using a quadrant position photodiode detector (Thorlabs) and steering the beam pointing with a piezoelectric actuated mirror (Thorlabs). A custom telescope, consisting of two achromatic doublets, upstream of the EOM precisely controls the beam diameter at the Powell lens.

A BBO electro-optic modulator (PCB4D-C-1030, Eksma Optics) rapidly (<10 ns) switches the beam between vertical and horizontal polarization states with high contrast, directing it into one or the other imaging path at a polarizing beam splitter. Only one path is illuminated at a time. In each path, the Fourier plane (pupil) of the line focus is imaged onto the surface of a custom 28-facet polygonal scanner (Lincoln Polygon), which scans each beam (*X*-axis). The use of a single scanner for the two paths reduces system complexity and ensures synchrony across paths. The polygonal scanner produces uniform pixel dwell times and unidirectional scans. A scan lens (SL50-CLS2, Thorlabs) forms the image of the scanned line, which is reimaged by a custom 1:1 relay onto the DMD surface.

SLAP2 uses DMDs designed for near-infrared light (DLP650LNIR, Texas Instruments; 800 ×1280 pixels,

10.8 µm pixel pitch, 10.8 kHz pattern rate, mechanical tilt angle ±12°), and a commercial FPGA-based controller (Vialux, V-650L). A single diffraction order (*m* = 3) of the DMD-modulated beam is relayed via a custom tube lens to the remote focus objective (Leica 11506529, 20 ×, NA 0.75, air), with a custom V-coating to maximize transmission at 1030 nm. The DMD is imaged onto the sample with an overall demagnification of 43.2 ×, so that each DMD pixel corresponds to 250 nm at the sample.

Each path’s three-dimensional imaging volume is 320× 200 × 450 µm (*x, y, z*). Each path’s z-axis is steered by a remote focus system (Botcherby et al. 2008). The lateral FOV is limited by the DMD pixel count and power requirements for generating the illumination line. In the image plane of each remote focus objective, a laser line mirror with linear piezo actuator (P-603, PI) performs independent axial (*Z*-axis) steering. The reflected, depth-steered beam returns through the same objective and custom tube lens, and having double-passed a quarter-wave plate, is directed through a PBS towards the front objective rather than returning toward the DMD.

The independently DMD-modulated and remote-focused beams each pass through a second custom tube lens, then are recombined at an image plane via a PBS. The overlapping beams pass through a final shared custom tube lens, and enter the sample-side objective (Leica HC IRAPO L 25× /1.00 W, 15507704; water immersion).

#### Detection path

As in conventional two-photon imaging, fluorescence emission is collected through the sample-side objective and directed by dichroic mirrors to a single-pixel detector for each color channel. Green-channel (iGluSnFR, ASAP7y) fluorescence is collected at 500–580 nm (Semrock FF01-540/80), and red-channel (jRGECO1a, RCaMP3) at 590–740 nm (Semrock FF01-665/150), separated by a 585-nm dichroic (Chroma T585lpxr). Customized silicon photomultiplier modules (SiPMs, Hamamatsu) are used for detection (Modi et al. 2019). A sliding primary dichroic directs visible light from the objective into either this two-photon detection path, or into an epifluorescence system during initial sample navigation.

#### DMD pattern generation

Different imaging modes can be combined within a single recording. To plan DMD pattern sequences, we decompose any set of user-specified ROIs into simultaneously-excited pixel sets (*superpixels*), then assemble DMD patterns one at a time using a greedy heuristic algorithm (`planAcquisition.m`) that prioritizes superpixels based on each ROI’s user-requested measurement rate. Two superpixels within the same column cannot be acquired in the same pattern. To avoid crosstalk due to the nonzero line width, superpixels within 8 columns of each other in the *X*-direction cannot be acquired in the same pattern, unless they lie within a threshold distance in the *Y*-direction. The planning algorithm iteratively (1) identifies superpixels that satisfy existing mutual exclusions; (2) prioritizes superpixels based on the ratio of their expected to desired measurement rates; then (3) greedily selects the longest run of highest-priority superpixels and adds it to the current pattern. When no superpixels can be added to the current pattern, another pattern is added. The sequence is complete when all superpixels are recorded at least once, and the imaging rate uniformity exceeds a threshold. 3D acquisitions are planned similarly, with additional constraints imposed by the dynamics of the remote focus actuator (`plan3DAcq.m`). The pattern generation algorithm and image formation process account for empirically-measured tilt or curvature of the illumination line at the DMD (“line shear”), calibrated via a user interface in the SLAP2 software. The *y*-resolution, numerical aperture, and power at the sample during raster imaging are set by the *Y dilation*, the number of contiguous rows of DMD pixels set to ON during each scan (an odd integer).

#### Data acquisition hardware and software

Full-featured commercial PC software for operating SLAP2 was developed by MBF Bioscience and was used for data acquisition. The SLAP2 graphical interface and control logic are implemented in MATLAB. A multithreaded C++ pipeline performs deconvolution, image generation, online motion correction, and streams resulting data to disk in an efficient proprietary format (.dat). Low-latency timing and safety operations are implemented by a field-programmable gate array (FPGA). A single PCIe DAQ card (MBF; 4-channel vDAQ) with an integrated FPGA performs data acquisition and input/output functions. Photodetector signals are digitized at 100 MHz, using a sample clock derived by 2× upsampling of the 50 MHz laser pulse train, so that digitization is phase-locked to the laser oscillator to avoid aliasing. SiPM single-photon pulses decay to 1% of their peak amplitude in ~200 ns, longer than the 20 ns laser period, so each digitized sample contains contributions from preceding laser pulses. The digitized signal is therefore deconvolved online by a fixed linear filter that outputs the number of detected photons phase-locked to each laser pulse, while suppressing non-phase-locked dark photons. The filter is fit using the instrument response function (IRF), which is estimated by weakly exciting a fluorescent sample with the laser, so that phase-locked single-photon events can be isolated and averaged.

#### Online motion correction

SLAP2 performs online motion correction by comparing the data from each acquired scan line, at ~ 10.8 kHz per path, to a dense 3D reference image (Figure S2). A running estimate of sample displacement is used to shift DMD patterns (*x,y*) and/or the remote focus position (*z*) to stabilize the image. The user can select any combination of pixel or band scanning ROIs within the field of view to be used for motion correction. Integration ROIs are not used for motion correction, but can be stabilized by including a separate pixel scan or band scan ROI within the FOV. A volumetric full-field raster stack acquired at the start of the recording serves as the 3D reference template. During random-access imaging, the acquired data are cross-correlated against the template, producing a normalized cross-correlation as a function of candidate x, y, and z displacements. Rather than performing the computation in a single pass at the end of an acquisition cycle, the correlation sums are accumulated incrementally as each line arrives (at the ~10.8 kHz line rate), so that at the end of the cycle the only computational steps that remain are normalization and finding a maximum. All template-dependent quantities that do not change between cycles—such as the template’s sum of squares and its normalization factor— are precomputed at the experiment start, so that the only per-line computation is accumulating the cross term between the incoming data and the template. The accumulation is vectorized with AVX2 fused-multiply-add instructions, and the template and correlation buffers are stored contiguously and ordered by memory address to maximize cache locality. The per-line update cost scales linearly (*O*(*N*)) with the number of superpixels *N* included in the motion estimate (~ 6 µs for a typical dendritic imaging experiment in this study), and the remaining end-of-cycle computation completes in <100 µs. The acquisition engine follows the peak of this cross-correlation objective function and updates the actuators: two registers on the FPGA-based DMD controllers shift the illumination pixel patterns laterally (x, y), and an analog voltage offset repositions each remote focus piezo axially (z). Because the last acquired line must still pass through the shared deconvolution and superpixel-formation pipeline before its contribution is added, the correction is issued ~0.2 ms after the end of data collection. After these updates are issued, the system waits for confirmation that they have taken effect by monitoring a digital trigger, whose change of state marks the end of the sync period; data acquired while the actuators are being updated are flagged as stale and excluded from the following motion estimate. Including this actuator-settling and sync interval, the correction is applied to the hardware ~0.7 ms after the end of data collection—small compared to the ~21 ms acquisition cycle. To minimize latency the sample is tracked with a small correlation search window; if the peak correlation falls below a settable threshold (i.e., lock is lost), the engine automatically widens to a larger cross-correlation search range to relocate the sample before reverting to the fast estimator.

### Optical characterization

#### Point spread function

Images for PSF characterization were collected using 200 nm orange fluorescent polystyrene micro-spheres (Lab 261) suspended in 1% or 3% agarose, covered with a cranial window to match *in vivo* conditions. Full-field raster stacks with 41 planes separated by 0.5 µm axially (20 µm volume height) were collected across the entire remote focus range. Each stack was averaged from 15 repeats, divided into 9 regions (3× 3 grid), and FWHMs were calculated from intensity profiles along the line of maximum integrated intensity aligned with each axis. The diffraction-limited numerical aperture (NA) was calculated from the z-FWHM averaged from all valid beads across depths (Zipfel et al. 2003).

#### Illumination uniformity

A single plane time series was collected using a yellow fluorescent slide (Thorlabs). To compute line uniformity, intensities were summed along the long (x) axis. Mean pixel intensities (normalized to unity) and standard deviations over time were calculated and plotted (Figure S1c). The illumination percentage variation across each DMD was calculated as the relative standard deviation of the mean pixel intensities.

#### Noise measurements

Pixel mean brightnesses and temporal variances (Figure S1d) were measured using a yellow fluorescent slide (Thorlabs) imaged at different laser powers. For these measurements, the scanner speed was adjusted to make each pixel an integer number of digitizer samples, avoiding effects of interpolation on image variances. The first 50 frames were discarded to eliminate thermal effects.

#### Field curvature

A transparent slide coated with a thin (<4 µm) layer of fluorescent dye was used to measure field curvature. A full-field raster stack with 41 planes separated by 1 µm axially was collected. For each unique xy pixel in the 3D volume, the z depth corresponding to the maximum intensity was located to form a maximum intensity surface. Curvature was fitted as a quadratic surface, detrended linearly to account for residual sample stage tilt.

### Simulations of SLAP2 imaging modes

#### Imaging speed comparison across methods

To compare imaging speeds of different microscopy methods *in silico*, we considered different numbers of sources uniformly distributed along dendritic branches, randomly distributed within a field of view. Synapse density was held constant at 0.5 synapse/µm, and ROI height was held constant across methods at 45 scan lines across the dendrite (5 bands in band scanning mode). For SLAP2, branch lengths were matched to the lengths of dendritic branch segments in our reference stacks. Branch orientation was not modeled explicitly: SLAP2 pixel and band scanning imaging rates depend only on the occupancy of DMD columns by ROI pixels. AOD acquisitions were simulated with slightly different parameters to reflect 3D drift scanning, for which imaging rates depend primarily on branch segment length, synapse density, and the number of scan lines across the dendrite. To reflect the ability of 3D AOD scanning to follow branches out of plane, AOD simulations used longer branch segments, with mean segment lengths derived from our reference volumes. For integration scanning, we assigned one integration ROI per synapse, and otherwise used the same simulation parameters as for pixel and band scanning. The simulated AOD parameters were based on TeO_2_ scanners (617 m/s sound velocity, ~55 resolvable spots/mm at 1030 nm), resulting in an 18.4 mm aperture for 1012 resolvable spots per axis and an access time of 29.8 µs. The number of resolvable spots was chosen so that the AOD system has the same number of addressable lateral positions (1012^2^≈ 1.02 ×10^6^) as a single SLAP2 path (1280× 800 DMD pixels). For simulated conventional raster scanning, we assumed a 12 kHz resonant scanner in bidirectional mode imaging the same total number of scan lines as SLAP2. The decrease in imaging speed as the number of imaged sources increases reflects the need to scan more lines to encompass those dendrites. The spatial density of dendrites in the simulation was selected to match those originating from a single neuron in our reference stacks, but (as for the other methods) the total number was extrapolated beyond the maximum we have observed in a single plane for Layer 2/3 cortical neurons. The dwell time per pixel for AOD and SLAP2 imaging was set to 36 ns, corresponding to SLAP2’s actual per-pixel dwell time for a polygon scan rate of 10.8 kHz. Simulations were repeated with 100 independent random samples for each number of simulated synapses plotted (1–1000).

#### Band scan parameter optimization

To determine the optimal width and spacing of bands used for band scanning, we simulated band scanning data at various dilations and subsampling factors. First, we collected the PSF of SLAP2 at multiple Y dilations. Then, we simulated two point sources aligned along the Y dimension, spaced 1 µm apart (Figure S4). A SLAP2 ROI 45 pixels tall in the Y dimension and 1 pixel wide in the X dimension was centered on these sources to image them. This ROI can have independently-tunable Y dilation and subsampling factor parameters. For each pair of Y dilation and subsampling factor, we subsampled along the empirical PSF for that Y dilation to get the simulated contribution of each of the two point sources to a superpixel sampling location. With the simulated imaging data from each point source, we quantified how distinguishable these two point sources were based on their SLAP2 imaging measurements. This was done by simulating 1000 noisy imaging samples for each point source, assuming Poisson noise with a photon count of 1 at the point source center, and calculating the Silhouette score of these 2000 samples between the two point sources (Rousseeuw 1987). As the metric could change depending on where along the vertical axis the samples were located, we varied the position of the two point sources (always keeping them 1 µm apart) and used the minimum Silhouette score across the different positions. We refer to this value as the “1 µm distinguishability.”

### Animals and surgery

#### Animal care and use statement

All experimental procedures involving animals were performed in accordance with protocols approved by the Institutional Animal Care and Use Committee at the Allen Institute (Protocol 2415), and conform to the National Institutes of Health (NIH) Guide for the Care and Use of Laboratory Animals.

#### Surgical procedures

C57BL/6J mice (Jackson Laboratory), regardless of sex, around two months of age, were implanted with a headplate and 5 mm diameter cranial window centered over primary visual cortex (Groblewski et al. 2020). Detailed subject, procedure, and session metadata are available alongside raw data released with this work. To achieve sparse labeling of neurons, mice were injected intracranially with a 150–250 nL mixture of AAVs encoding reporters and a low titer (1.2E8–6E8 GC/mL) of AAV9- or AAV-PHP.eB-CaMKII0.4-Cre (pENN.AAV.CamKII 0.4.Cre.SV40, Addgene: 105558) in the visual cortex (approx. − 3.9 mm A/P, −3.1 mm M/L from Bregma). For glutamate imaging, 4E12–5E12 GC/mL AAV-PHP.eB-hSyn-FLEX-iGluSnFR4f (Addgene: 234442) or 5E12 GC/mL AAV-PHP.eB-hSyn-FLEX-iGluSnFR4s (Addgene: 234441) was used. For calcium imaging, 1E12 GC/mL AAV-PHP.eB-CAG-FLEX-jRGECO1a (Addgene: 100852) or 2E12–5E12 GC/mL AAV-PHP.eB-CAG-FLEX-RCaMP3 (Yokoyama et al. 2024) was used. For voltage imaging, 1E12 GC/mL AAV-PHP.eB-hSyn-DIO-ASAP7y (Hao et al. 2026) was used. Mice were imaged at least 6 weeks after viral injection and headplate implantation.

### Imaging experiments

#### Visual stimulation

Visual stimuli were presented via an OLED screen placed 15 cm from the mouse’s right eye. Stimuli were presented using only the blue channel of the screen, with an additional optical filter (Wratten 47B) to prevent stimulus light bleedthrough into detection channels. Moving grating visual stimuli (2 Hz temporal frequency, 0.04 cycles/deg) of 8 directions (45-degree increments) and a blank screen of mean luminance were presented in randomized order, each for 2 s and spaced by 1 s of mean luminance. Each stimulus was presented at least 40 times in glutamate recordings and 20 times in voltage recordings. Stimuli were implemented using the BonVision package within the Bonsai programming language (Lopes et al. 2021; Lopes et al. 2015), with behavioral video and synchronization signals recorded using a Harp Behavior device (Carvalho et al. 2024).

#### Pixel scanning

For pixel scanning experiments, neurons of interest expressing iGluSnFR4f and jRGECO1a (or RCaMP3) were first identified. One plane was selected on each imaging path, one containing the soma and perisomatic dendrites and one containing basal or apical dendrites, and a reference stack (21 planes, 1 µm apart) was collected around each plane using SLAP2 full-field raster imaging. SLAP2 imaging ROIs were drawn around the soma and the dendrites belonging to that soma and were set up for pixel scanning with a Y dilation of 5–7. Excitation power was controlled by the half-wave plate, which was set to 70–100% for these experiments (20–70 mW at the sample). The pixel-scanning dataset consisted of 20 sessions from 7 mice. One neuron was imaged in two sessions, in which one of the two imaging planes was different.

#### Interleaved pixel and band scanning

We conducted interleaved pixel and band scanning experiments of the same dendrites to compare the data between the two imaging modes. In these experiments, a plane with dendrites was found on each imaging path and a reference stack (at least 20 µm thick) was collected around each plane. Imaging ROIs containing iGluSnFR4s-expressing dendrites were drawn on a single plane in each path and were set up to do both band scanning (subsampling factor of 9 and Y dilation of 17) and pixel scanning. With these settings, the SLAP2 acquisition alternated between pixel scan superpixels and band scan superpixels, so that within a cycle, each full set was acquired at least once. Band scan superpixels were sampled at an average of 212 Hz and pixel scan superpixels were sampled at an average of 106–423 Hz. Excitation power was controlled by the half-wave plate, which was set to 70% for these experiments. The interleaved pixel and band scanning dataset consisted of 4 imaging planes across 2 mice.

#### Volumetric band scanning

For volumetric band scanning experiments, neurons of interest expressing iGluSnFR4s and RCaMP3 were first identified and reference stacks capturing their dendritic arbors were collected using SLAP2 full-field raster imaging up to a day before the experiments. After collecting the reference stack and before the experiment, SLAP2 imaging ROIs were selected across several planes in this volume to identify a set of dendrites belonging to the targeted neuron. An ROI was also drawn around the soma. The number and arrangement of ROIs was selected to ensure a volume rate on each imaging path of at least 50 Hz. All ROIs were set up for band scanning with a subsampling factor of 9 and Y dilation of 17. The band scanning dataset consisted of 6 sessions of 6 different neurons from 3 mice.

#### Integration scan voltage imaging

For integration scanning experiments, neurons of interest expressing ASAP7y were first identified. One plane was selected on each imaging path, one containing the soma and perisomatic dendrites and one containing apical dendrites, and a reference stack (21 planes, 1 µm apart) was collected around each plane using SLAP2 full-field raster imaging. SLAP2 imaging ROIs were drawn around the soma and dendritic segments belonging to that soma and were set up for integration scanning at a sampling rate of 2 kHz per ROI. ROIs were refined using an automated script to restrict illumination to labeled pixels with a small surrounding buffer region. One or more pixel scan ROIs imaged at >200 Hz were included for real-time motion correction (see **Integration scanning: trace extraction**). Excitation power was controlled by the half-wave plate, which was set to 65–80% for these experiments (<70 µW/µm^2^). The voltage imaging dataset consisted of 22 sessions from 5 mice.

### Preprocessing

#### Image generation

Within each cycle through the DMD pattern sequence, a given superpixel can be measured multiple times, and different pixels are measured at different times. We interpolate these irregular measurements onto a regular query timebase using exponential time weighting (e.g. time constant 5 ms for 200 Hz query timebase). Throughout analysis, we propagate estimates of both the superpixel brightness and an associated uncertainty factor (*freshness*) that tracks the varying Poisson noise associated with each superpixel intensity estimate and depends on the number and time weights of the contributing measurements.

#### Pixel scanning: post-hoc motion correction

Images were generated with an 80 Hz query timebase and aligned using a custom algorithm (MultiRoiRegistration), adapted from StripRegistration (Xie et al. 2026) to accommodate thin, sparsely sampled fields of view. In brief, the algorithm first initialized a template by aligning a subset of mutually well-correlated frames from 40 evenly-spaced frames throughout a trial. Each frame of the movie was then aligned against the template by finding the X and Y displacement that yielded the maximum freshness-weighted normalized cross-correlation among imaged pixels. The pixel values and template values were square root transformed to stabilize variance. Subpixel shifts were determined by fitting a quadratic along each dimension to the cross-correlation function around the maximum. The peaks of the quadratic were selected as the motion displacement estimate in each direction. With the displacement estimate, the motion corrected frame was generated by linear interpolation, weighted by the freshness of each contributing observation. The motion corrected frame was then averaged into the template to dynamically update it as frames become aligned. In cases where a Z motion estimate was desired, the dynamic template was not used. Instead, we used the full-field raster scanned reference volume as a 3-dimensional template for alignment.

#### Pixel scanning: source extraction

Source extraction for pixel scan data was performed using Source Identification by Activity Localization (SILo), implemented in MATLAB (Xie et al. 2026). In brief, the method takes inspiration from superresolution localization microscopy methods (Lelek et al. 2021; T.-W. Chen et al. 2025) to precisely identify source locations despite the reduced effective resolution produced by the pixel size. We modeled a single event as having a spatiotemporal profile of a small Gaussian dot (standard deviation of 1.33 pixels) modulated over time by a decaying exponential of a time constant matched to the glutamate indicator. We performed event detection by convolving the reflection of this shape with a background-subtracted, noise-normalized version of the movie (sampled at the same 80 Hz timebase used for motion correction) to identify local maxima in space and time (i.e., a 3D matched filter). These events were weighted by their intensity in the filtered movie and aggregated in quadrature over time into a summary image, which we term the *activity image*. The activity image shows localized densities around active sources. We then identified the centroids of these densities by fitting each to a symmetric Gaussian to establish the location of each synapse. As a final step, we used the established source locations as the initialization for constrained non-negative matrix factorization on the recording data sampled at a 200 Hz timebase to fine tune the spatial profiles and extract their corresponding time traces. The matrix factorization provided a denoised estimate of Δ*F* for each source and an estimate of the background movie. The *F*_0_ for each source was then calculated as the background movie projected onto each spatial footprint, normalized by its squared norm. We also calculated an unconstrained least-squares fit of the set of spatial profiles onto the background-subtracted data to get a “least squares trace” corresponding to each source. This trace was used for characterization of noise in the source traces.

Somatic calcium activity was extracted by spatial averaging of raw movie pixels within a manually selected polygon corresponding to the soma. The baseline fluorescence *F*_0_ of the somatic calcium trace was estimated as a slowly varying lower envelope of the trace itself: the trace was median filtered (1 s window), a rolling minimum was taken over a 4 s window on a decimated time grid, samples next to missing values were discarded, and the envelope was smoothed and interpolated back to the original sampling by piecewise cubic Hermite interpolation. Δ*F*/*F* was computed as (*F* − *F*_0_)/*F*_0_.

#### Band scanning: post-hoc motion estimation

For estimating and correcting residual motion post-hoc, band scan imaging data were queried at a timebase with a specified frequency (30 Hz for volumetric band scan recordings and 80 Hz for interleaved pixel and band scan recordings). Residual motion offsets were estimated by comparing data to a reference stack collected at the same Y dilation as the band scans. At each time step, the optimal motion displacement was the three-dimensional translation that maximized the cross-correlation of the data with the reference stack. Similar to pixel scan data, subpixel shifts were determined by fitting quadratics along each dimension to the cross-correlation function around the maximum and taking the peaks. Post-hoc motion offsets estimated from band scan data alone were within 0.14 µm of those estimated from pixel scan data in recordings with interleaved modes (90th percentile; Figure S8b).

#### Band scanning: source extraction

Source extraction for band scan data was performed using BandSILo, a variant of the SILo algorithm (Xie et al. 2026) used for pixel scan data, and was implemented in Python. Similar to SILo, BandSILo proceeds in two major steps—activity image generation and trace extraction—but also accounts for the subsampled band geometry and residual sample motion within these steps.

Activity image generation: First the data was queried on a timebase matching that used for motion correction. The background fluorescence was then estimated by interpolating the subsampled band scan data—shifted by the corresponding post-hoc motion estimate—in each frame onto the dense pixel grid using natural cubic splines. A temporal rolling median (over a 4 s window) was calculated on the dense grid, which was then subsampled back into band scan superpixel (accounting for motion) to get the background fluorescence of the band scan data. The background-subtracted data was then normalized by an estimate of the noise at each superpixel, using the method described in Xie et al. 2026. Glutamate transients were then identified on this normalized data by a spatial matched filter corresponding to a center-surround profile based on the point spread function for the band scan’s Y dilation and a temporal matched filter of an exponential decay in time (time constant: 150 ms). Local maxima in this movie were aggregated over time in quadrature and spatially high-pass filtered to generate an activity image.

Trace extraction: Source centers were initialized by the peaks of Gaussians fit to the activity image. The source spatial profiles were then refined by a non-negative matrix factorization of the background-subtracted, normalized data, with spatial profiles constrained to Gaussians convolved with the band scan PSF. Sources with traces that had low signal-to-noise ratio were discarded. With these spatial profiles, the final traces were extracted from band scan data that was queried at a 50 Hz timebase. This data was background-subtracted using an upsampled version of the background data from above. Denoised Δ*F* traces for all sources were fit by a constrained least squares fit to this data with the spatial profiles held fixed. Denoised traces were constrained to be nonnegative event amplitudes convolved with a 150 ms exponential decay kernel. An L1 sparsity penalty was also applied to isolate the sparse events. In addition to the denoised trace, a least-squares trace was also extracted by an unconstrained least-squares fit of all the sources to the background-subtracted data.

Somatic calcium extraction: Somatic calcium activity was extracted by spatial averaging of raw superpixels within a manually selected polygon corresponding to the soma. *F*_0_ and Δ*F*/*F* were computed in the same way as for pixel scanning.

#### Integration scanning: trace extraction

Voltage imaging recordings included one or more pixel scan ROIs imaged at >200 Hz for real-time motion correction. We first performed post-hoc motion correction using pixel information to identify residual motion, and censored all frames showing uncorrected displacements of more than 500 nm. Detected photon rates for each ROI were interpolated onto a regular time grid using exponential time averaging (time constant 16 line periods, ~1.48 ms). Δ*F*/*F*_0_ was computed per trial with *F*_0_ estimated by low-pass filtering *F* (zero-phase second-order Butterworth filter, 0.1 Hz cutoff).

#### Signal-to-noise ratio (SNR) estimation

Noise was estimated as the standard deviation of the derivative of the activity traces divided by 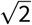. Signal was estimated as the 95th percentile of peak amplitudes, detected using the `find_peaks` function from the SciPy Python library. SNR for each trace was calculated by its signal measurement divided by its noise estimate.

#### Estimation of synapse density

Synapse density was defined as the total number of imaged synapses divided by total imaged dendritic length. Dendritic length was estimated by binarizing the mean fluorescence image via local adaptive thresholding, skeletonizing each connected dendrite segment, and summing geodesic skeleton lengths across both imaging paths. For band scanning data, instead of using the mean fluorescence image, the reference stack masked to only the pixels within imaging ROIs was used to estimate dendritic length. The soma (manually annotated) was excluded prior to length estimation in both cases.

#### Neuron tracing

Neurons were traced using the SNT ImageJ plugin (Arshadi et al. 2021). We discarded any imaged synapses from functional recordings where the centroid of the ROI was > 5 µm from the nearest traced dendrite.

### Characterization of glutamate inputs and calcium output

#### Identification of stimulus-modulated synapses and somata

We identified stimulus-modulated synapses by quantifying the reliability of their responses across trials. For each synapse, we computed the time-resolved mean peri-stimulus response per stimulus identity (the synapse’s *tuning profile*). We computed a cross-validated measure of trial-to-trial similarity by correlating each trial with the mean response of the remaining trials (leave-one-out). To assess significance, this reliability metric was compared to a null distribution obtained by independently shuffling the time points of each trial (500 permutations), which preserves overall activity statistics while disrupting alignment to the stimulus sequence. *P* values were computed as (*n*_null *≥*obs_ +1)/(500+1) and corrected for multiple comparisons with the Benjamini–Hochberg false discovery rate (*q* < 0.01), across the synapses of each session for synapses and across sessions for somata. ROIs passing this threshold were classified as stimulus-modulated. Somata were tested in the same way.

#### Cross-correlation of somatic and summed synaptic activity

For each session, the somatic Ca^2+^ signal and the summed glutamate input were cross-correlated trial by trial (lags up to ±5 s). The lag of the session was the peak of the trial-averaged cross-correlation, on a 5 ms grid; negative lags indicate that the input leads the soma. Lags outside 1.5 × IQR were excluded as outliers (3 of 20 sessions), and the mean ± SEM of the remaining 17 sessions is reported.

#### Orientation and direction selectivity

The orientation selectivity index (OSI) was computed from the baseline-corrected, trial-averaged responses across all eight stimulus directions (0°, 45°, …, 315°) using the vector sum method:

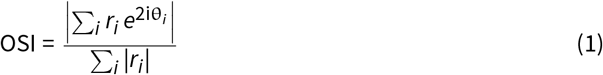

where *r*_*i*_ is the mean response during the 2-second stimulus presentation window for direction *i*, and θ*i* is the corresponding stimulus orientation in radians.

The preferred orientation was defined as 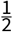 arg 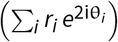 and mapped to the range [0°, 180°).

The direction selectivity index (DSI) and preferred direction were computed in the same way from the vector sum without doubling the angle,

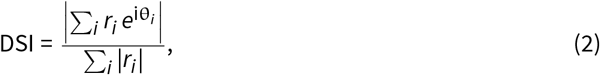

with the preferred direction defined as arg 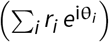 and mapped to the range [0°, 360°).

Analyses on pixel scanning data based on preferred orientation were restricted to stimulus-modulated sessions with soma OSI > 0.3 (15 of 17) and, for synapses, synaptic OSI > 0.3.

#### Non-negative matrix factorization of stimulus responses

NMF was performed on a 3-second window of trial-averaged peri-stimulus activity, for all synapses pooled across 20 sessions to each of the presented stimuli; i.e. the factorized matrix had dimensions [33325 (synapse × stimulus; 8 grating directions plus blank screen where available) × 600 (time-points)]. Peri-stimulus activity was computed as for tuning profiles. Δ*F*/*F* responses were used with-out further normalization, and negative values were clipped to zero. NMF was implemented using `sklearn.decomposition.NMF` in Python, with `n_components=10`. The fraction of signal energy captured by the factors was quantified as

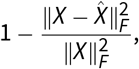

where *X* is the original data matrix, 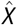 is its NMF reconstruction and ∥·∥_*F*_ is the Frobenius norm (the square root of the sum of squared entries). The factors captured > 94% of the signal energy of *X*. We grouped the resulting components into 4 major categories (ON (2 components), OFF (2 components), CYC (4 components), SUP (1)), and a DC offset. Aggregated weights reported for these categories are the mean of the weights of the raw components. For each synapse, the weight of a category was summed across stimuli, and the *proportion* of a category was this weight divided by the summed weight of the four categories (ON, OFF, CYC and SUP; the DC offset was excluded), so that the four proportions of a synapse sum to 1. To compare NMF weights of inputs and outputs, we applied the fitted NMF model to deconvolved calcium tuning curves. Deconvolution was performed using OASIS (Friedrich et al. 2017) to correct for indicator dynamics. Correlation analysis within each component group used the Pearson correlation coefficient between 8-element direction-weight vectors.

#### GLM-based simulations of neuronal output from synaptic inputs

To simulate somatic output from synaptic input, we assembled a set of synaptic tuning profiles by pooling across all sessions (3826 synapses across 20 sessions). Because individual recordings varied in depth of soma and dendritic plane as well as location in V1, we treated the pooled set as representative of the input types received by L2/3 neurons. For each simulated neuron, we drew *N* = 3,000 synapses from this pool by bootstrap (sampling with replacement). Individual trials were generated by adding three Gaussian terms to the tuning profiles: a multiplicative gain shared across all synapses, an independent multiplicative gain per synapse, and additive noise per synapse with a standard deviation of 0.1 times the mean absolute response of the pooled synapses.

For direction-amplifying neurons, synaptic weights were drawn from a Gaussian prior with mean proportional to a von Mises function of the difference between each synapse’s preferred direction and the soma’s target preferred direction (exp(κ cos(θ_soma_ −θ_*i*_)), where κ sets the strength of directional tuning), scaled by the synapse’s direction selectivity index. The prior had a standard deviation of 1, and weights were normalized to sum to 1; the same prior width and normalization were used for ON-amplifying neurons.

For simulations amplifying the ON temporal category, synaptic weights were drawn from a Gaussian prior with mean proportional to an exponential function of each synapse’s ON proportion (exp(κ_on_ ·*f*_ON_), where κ_on_ sets the strength of ON amplification).

Somatic activity was generated as a weighted sum of single-trial synaptic traces, standardized, passed through an exponential nonlinearity with temperature τ (exp(*z*/τ)) and scaled to its 99.5th percentile, defining the ground-truth soma.

To model experimental condition, we subsampled 5% of synapses at random and added independent Gaussian measurement noise to both the observed synaptic traces and the soma, with standard deviation equal to the mean absolute signal divided by the signal-to-noise ratio.

Across simulated datasets, we varied the soma’s preferred direction (drawn uniformly from the eight stimulus directions), the standard deviations of the shared and independent multiplicative singletrial gains (*N* (0.1, 0.1), minimum 0.01, and *N* (0.3, 0.3), minimum 0.05), κ (directional tuning strength; *N* (4, 2), minimum 0.5), the exponent of the direction selectivity index in the weight prior (*N* (2, 1), minimum 1), and κ_on_ (ON-factor strength; *N* (6, 2), minimum 2). The temperature of the exponential nonlinearity was drawn from *N* (0.75, 0.25) (minimum 0.5) for direction-amplifying neurons and from *N* (0.5, 0.15) (minimum 0.3) for ON-amplifying neurons, and the signal-to-noise ratio of the soma from *N* (5, 3) (minimum 1); observed synapses had a signal-to-noise ratio of 10. Here *N* (µ, σ) denotes a normal distribution with mean µ and standard deviation σ.

The example simulations shown in Figure S5g–h used κ = 4, a direction selectivity exponent of 3 and a temperature of 1.25 (direction-amplified example), and κ_on_ = 4 and a temperature of 1 (ON-amplified example), with standard deviations of 0.1 and 0.3 for the shared and independent single-trial gains and a somatic signal-to-noise ratio of 2. Simulated datasets in which the unweighted summed input explained a larger fraction of somatic variance than in any observed session, and which may therefore have been easier to fit, were excluded from the summary panels (5 of 24 simulations); including them did not change the result qualitatively (optimally weighted *R*^2^ of 0.82–0.99 for all 24 simulations).

#### Activation functions

Optimally weighted activation functions were produced by fitting a single exponential GLM (soma≈ exp(β_0_ + Σ_*i*_ *w*_*i*_ ·synapse_*i*_)) that assigns one weight per synapse, then using those weights to compute a weighted sum of synaptic input. The moment-to-moment somatic output (Δ*F*/*F*) was then plotted against this GLM-weighted summed input (Δ*F*/*F*) to obtain the activation function. Weights are termed *optimal* because the model was fit to the entire recording without splitting trials into train and test sets; an in-sample fit gives an upper bound on how much of the somatic output synaptic weighting can explain, so a low explained variance is a conservative result. The resulting weights therefore represent the best estimate from the data, rather than an estimate validated on held-out trials.

Activation functions were similarly computed for each NMF temporal factor category. Using the fitted NMF model, the full trace for each synapse and the soma was decomposed into the four factor categories (CYC, ON, OFF, SUP), and the same per-synapse weights described above were used to compute the weighted summed input for each category. Somatic output was then plotted against this summed input per category. Residuals were defined as the observed somatic trace minus the prediction of the fitted single-predictor exponential activation curve, evaluated separately for the weighted and unweighted summed input. Residuals were summarized either per condition (mean residual per stimulus direction, relative to preferred direction, or per NMF factor category) or as a residual time course (mean residual across conditions, resolved within trial time).

### Spatiotemporal organization of inputs

#### Distance-dependent analyses of synaptic tuning

Cross-validated stimulus correlations between pairs of synapses were computed using a split-half procedure. Trials were randomly divided into two groups, tuning profiles calculated for each split, and the Pearson *r* computed between the tuning profile of one synapse in one half and that of the other synapse in the other half. This was repeated 100 times with different random splits and averaged. Cross-validated correlations between somatic output and synapses used 200 random splits and accounted for indicator dynamics by taking the signed correlation at the lag between 0 and 200 ms with the largest absolute value.

Noise correlations were computed as the Pearson correlation between trial-to-trial residuals after regressing out tuning profiles (with each trial’s tuning profile computed from remaining trials), after smoothing the traces with a 20 ms moving average. Noise correlations between synapses and the soma were the signed values at the lag between 0 and 200 ms with the largest absolute value.

The relationship between correlation strength and distance was tested using ANCOVA with the correlation as the dependent variable, distance as a continuous predictor, and session ID as a categorical factor, including their interaction. The ANCOVA model was implemented using `statsmodels` in Python.

The dependence of the proportion of each temporal factor category on Euclidean distance to the soma was tested with an ANCOVA, with distance as a continuous predictor, session as a categorical factor and their interaction. To compare dendritic regions, synapses were classified by their depth relative to the soma: synapses more than 5 µm above the soma were classified as apical and all others as basal. The difference between regions was tested for each proportion with a two-way ANOVA, with region and session as categorical factors and their interaction, using only sessions with synapses in both regions. Both tests were implemented using `statsmodels` in Python (Type II sums of squares), and the reported *F* and *p* values are for the distance or region term.

#### Noise correlation tuning curves

Trial-to-trial noise was isolated by regressing out the tuning profile from each synapse and soma trace using leave-one-out mean regression. Time-resolved pairwise noise covariance was computed across trials at each time point for all synapse–soma and synapse–synapse pairs. Population-averaged noise covariance traces were normalized by a single scalar: synapse–soma covariance by the product of the soma standard deviation and the mean synapse standard deviation across the population; synapse–synapse covariance by the mean synapse variance across the population.

### Predictive modeling of input-output computations

We used a generalized linear model (GLM) to predict somatic output from synaptic input in both pixel scan and volumetric band scan datasets. At each time point *t*, the predicted somatic signal *ŷ*_*t*_ was expressed as an exponential function of the sum over temporally filtered synaptic inputs together with stimulus and timing information:

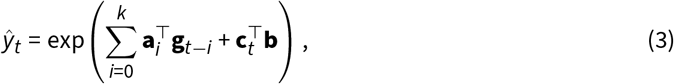

where **g**_*t −i*_ represents the synaptic input at time lag *i*, **a**_*i*_ are temporal filter coefficients, **c**_*t*_ is a one-hot encoded vector representing the stimulus and timing condition at time *t*, and **b** is the corresponding weight vector. Model parameters were estimated by maximizing the likelihood of the somatic signal under a Gamma distribution. The measured somatic signal used was a version of the Δ*F*/*F* trace that was denoised using OASIS (Friedrich et al. 2017).

Parameter estimation was performed by minimizing the negative log-likelihood with an *L*_2_ regularization term. Trials were split into five train-test splits, with non-overlapping test-trial groups across splits, and results were aggregated across the five splits. For each split, 20% of trials were held out as the test set. The regularization strength and temporal filter length were optimized using cross-validation within the training set. When allowing synaptic weights to vary, weights were initialized using uniform weights derived from the corresponding total input model, and fitted weights were rejected if they performed worse than the total input initialization on the validation set. The constrained model was fit using `glum` in Python, and other models were fit using `scikit-learn`.

#### Model variants

To compare with previous experimental paradigms and assess how different forms of synaptic input contribute to somatic output, three GLM variants were fit.

Total input model: The summed synaptic activity across all synapses in each trial was used to predict the somatic signal on a single-trial basis. This captures trial-to-trial fluctuations in total synaptic drive, but does not resolve individual synapses. It represents a model that could be fit from integrative measurements that lack single-synapse resolution, such as photometry or conventional widefield imaging. The single-trial total input model is

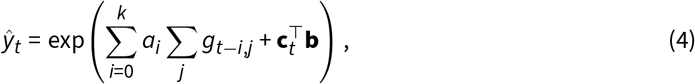

where *a*_*i*_ is the weight for lag *i, g*_*t−i,j*_ is the activity of synapse *j* at time *t* − *i*.

Subsampled-synapse model: Data from different numbers of randomly selected synapses were used as predictors for a series of models. Each model was trained and tested with single-trial data from the selected synapses. These models were used to assess how the number of simultaneously measured synapses affects prediction of somatic output. The subsampled-synapse model is

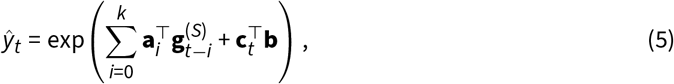

where 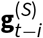 is the synaptic activity restricted to a subset *S* of all synapses, **a**_*i*_ is the weight vector for lag *i*.

Model with a nonnegative exponentially decaying kernel: To better interpret the GLM weights, we fit a variant in which synaptic inputs were combined temporally using an exponentially decaying kernel, with the kernel weights constrained to be nonnegative. The model is

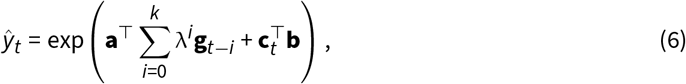

where λ is the decay factor, and **a** is the weight vector applied to the exponentially weighted sum of synaptic inputs.

### Analysis of interleaved pixel and band scan experiments

Data preprocessing (pixel scan and band scan), SNR calculation, and OSI calculation were performed in the same manner as described above. In a recording, pixel scan sources and band scan sources were considered matches if their denoised traces had a correlation of at least 0.1 and their spatial profile centers were within 1 µm along the *x*-axis and 1.5 µm along the *y*-axis of each other. The matching was more lenient along the *y*-axis as band scanning has lower resolution along the *y*-axis, leading to more uncertainty in location estimation along that direction. To identify one-to-one matches, a maximum bipartite matching algorithm was used, where each match by the trace and spatial profile criteria denoted an edge between sources.

### Voltage data and oscillation analyses

#### Correlation and lag between dendritic compartments

Correlations and lags between compartments (Figure 6d) were computed on the Δ*F*/*F*_0_ traces, which were exponentially time-averaged onto a regular time grid at the line rate (~10.8 kHz; see **Integration scanning: trace extraction**), without further smoothing or downsampling. For each trial and each pair of compartments we computed the Pearson correlation and the normalized cross-correlation of the mean-subtracted traces; the lag of a pair was the position of the largest absolute cross-correlation within ±0.5 s. Correlations were averaged over trials and then over all pairs of the same type (soma– apical, soma–basal, apical–basal) within a session; the lag of a session was the median over all trials and pairs of the same type. Positive lags indicate that the second compartment of the pair lags the first. Significance was assessed with a two-sided Wilcoxon signed-rank test against zero across sessions.

#### Detection of oscillatory events

In glutamate recordings, oscillations were detected from summed glutamate inputs; in voltage recordings from activity in the somatic compartment. Detection was performed by identifying localized events in time-frequency spectrograms (2.5–10 Hz) using the `blob_dog` function from scikit-image. Peak time and frequency were refined by sub-resolution localization of the spectrogram maximum. Oscillation start and end times were estimated by tracking contiguous regions in the spectrogram surrounding each peak.

#### Somatic calcium response to glutamate oscillations

Somatic Ca^2+^ and total glutamate Δ*F*/*F* were aligned to the glutamate peak nearest to each detected oscillation peak (within 0.25 s). Each event trace (± 0.5 s) was mean-centered and averaged per session. To test for modulation of somatic calcium, the mean in each of two windows after the peak (0–0.25 s and 0.25–0.5 s) was compared with a baseline window (− 0.5 to −0.25 s) across sessions using a paired Wilcoxon signed-rank test.

#### Synapse activation during glutamate oscillations

For each session, glutamate traces from individual synapses and the total glutamate signal (mean across synapses) were band-pass filtered at 2.5–6.6 Hz, corresponding to the 95% interval of glutamate oscillation frequencies. For each trace, segments corresponding to detected oscillatory events (from event onset to offset) were extracted and concatenated end-to-end into a single trace. Synapse activation was then quantified as the Pearson correlation between the concatenated trace of each synapse and the corresponding concatenated total glutamate trace, resulting in a single activation value for each synapse per session.

**Figure S1:**
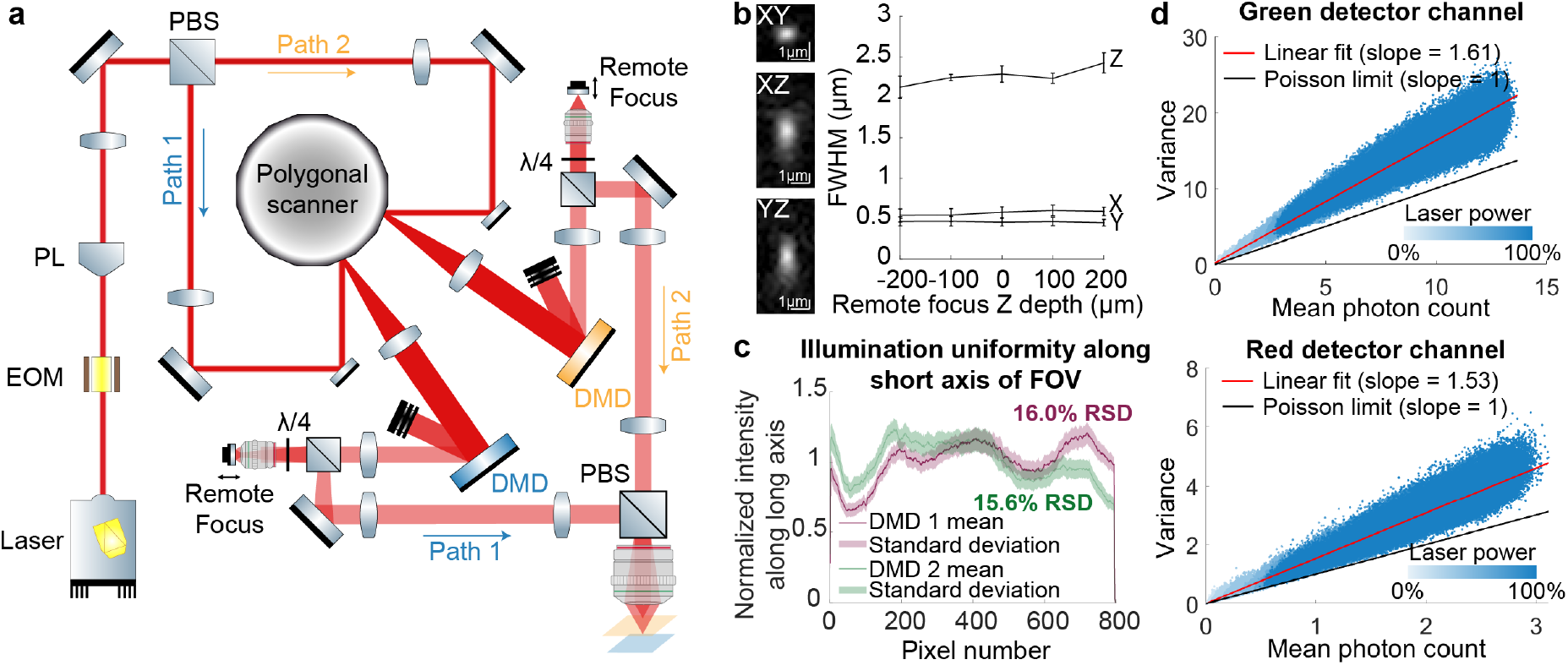
SLAP2 optical design and properties, related to Figure 1. **a**, SLAP2 optical schematic (simplified). A powerful laser is shaped into a high-aspect-ratio line focus by a Powell lens, and switched between two imaging paths using an EOM and PBS. For each path, the polygonal scanner repeatedly scans the line across a DMD, which programmably shapes the illumination light. Independent remote focus systems steer each path in depth. The two paths are recombined at a PBS and directed to the sample. Fluorescence is detected with single-pixel detectors (not shown) as in conventional two-photon imaging. **b**, Measured PSF (200 nm fluorescent bead) at remote focus 0 µm, as XY XZ and YZ slices through the PSF. Images are intensity-normalized with gamma = 0.5. (Right) Full-width half maxima of PSFs along each axis (0.57*±* 0.07 µm in X, 0.46*±* 0.05 µm in Y, 2.27*±* 0.14 µm in Z) recorded across remote focus depths. **c**, SLAP2 illumination uniformity along the short axis of each DMD. RSD: relative standard deviation of mean pixel intensities along the short axis. **d**, Detector noise at various total laser powers and linear fit of green and red channels. The Poisson limit (slope = 1) corresponds to a perfect detector. The SLAP2 detectors mainly exhibited multiplicative noise due to pulse height variability (Modi et al. 2019).

**Figure S2:**
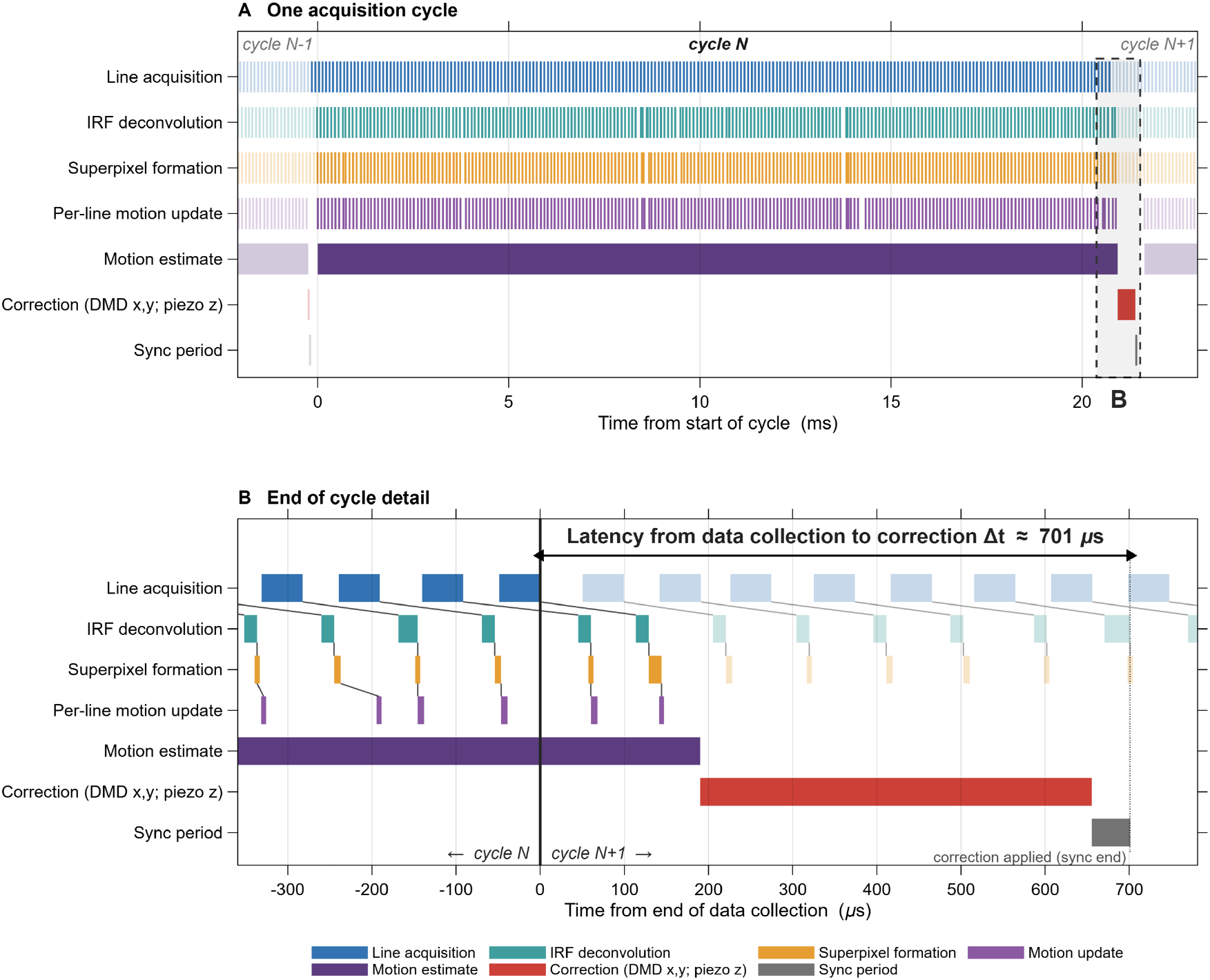
Real-time data flow and latency of the SLAP2 online motion-correction pipeline, related to Figure 1. Timeline representation of the multithreaded C++ pipeline during *in vivo* pixel imaging. Each row is a stage in the processing pipeline; horizontal bars mark the start and end of each operation, so data flow proceeds top-to-bottom (acquisition → processing → correction) and left-to-right (time). **a**, One full acquisition cycle (cycle N; 224 lines, ~20.9 ms; ~10.8 kHz line rate). Successive lines stream continuously through the pipeline — line acquisition → IRF (instrument response function) deconvolution → superpixel formation → per-line motion update — while the motion estimate (a sparse normalized cross-correlation against the reference volume) is accumulated incrementally over the whole cycle. At the end of the cycle the resulting displacement is issued as a single correction (lateral DMD x,y offset and axial remote focus piezo z) and applied to the hardware (sync period). Line-acquisition bars show only the DMD active illumination window (~ 50% of the line period), during which this path collects light. The pipeline for only one of the two interleaved DMD paths is shown; the remainder of each line period is used to acquire the second DMD path, which runs an identical independent pipeline (not shown). The central cycle (cycle N) is drawn opaque, with the preceding (cycle N-1) and following (cycle N+1) cycles shown semi-transparent to indicate that the process repeats continuously. The dashed box marks the region expanded in (b). **b**, End-of-cycle detail, showing the latency between the end of data collection and the applied correction. Thin black connectors trace each individual line scan through its processing stages (acquisition → deconvolution → formation → motion update). t = 0 is defined as the end of data collection (the last DMD sample contributing to the estimate); the vertical divider separates cycle N (opaque) from cycle N+1 (semi-transparent), whose acquisition has already begun. The motion estimate is finalized ~0.2 ms after the last sample, the correction is written to the DMD registers and remote focus piezo, and the sync period ends when a digital trigger from the DMD’s FPGA-based controller confirms the update has taken effect — giving an end-to-end latency of Δ*t* = 0.7 ms from the end of data collection to the applied correction, compared to the ~21 ms cycle period. Data acquired during this latency period is considered stale and not used for the next correction.

**Figure S3:**
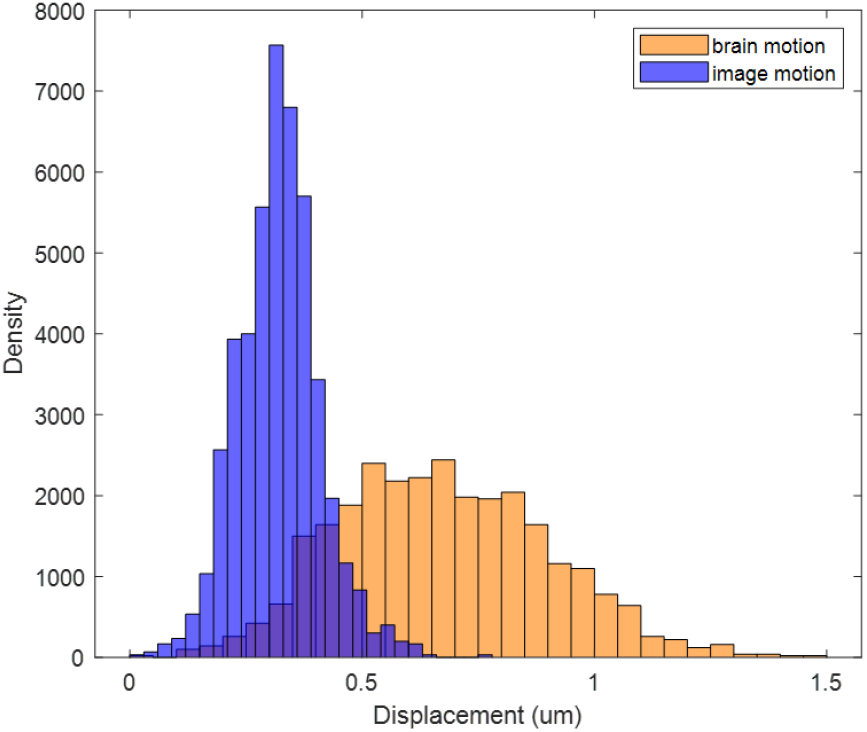
Online motion correction performance during band scanning, related to Figure 1. Distribution of brain motion (orange) and residual image motion (blue) during a band scanning acquisition on SLAP2 with online motion correction.

**Figure S4:**
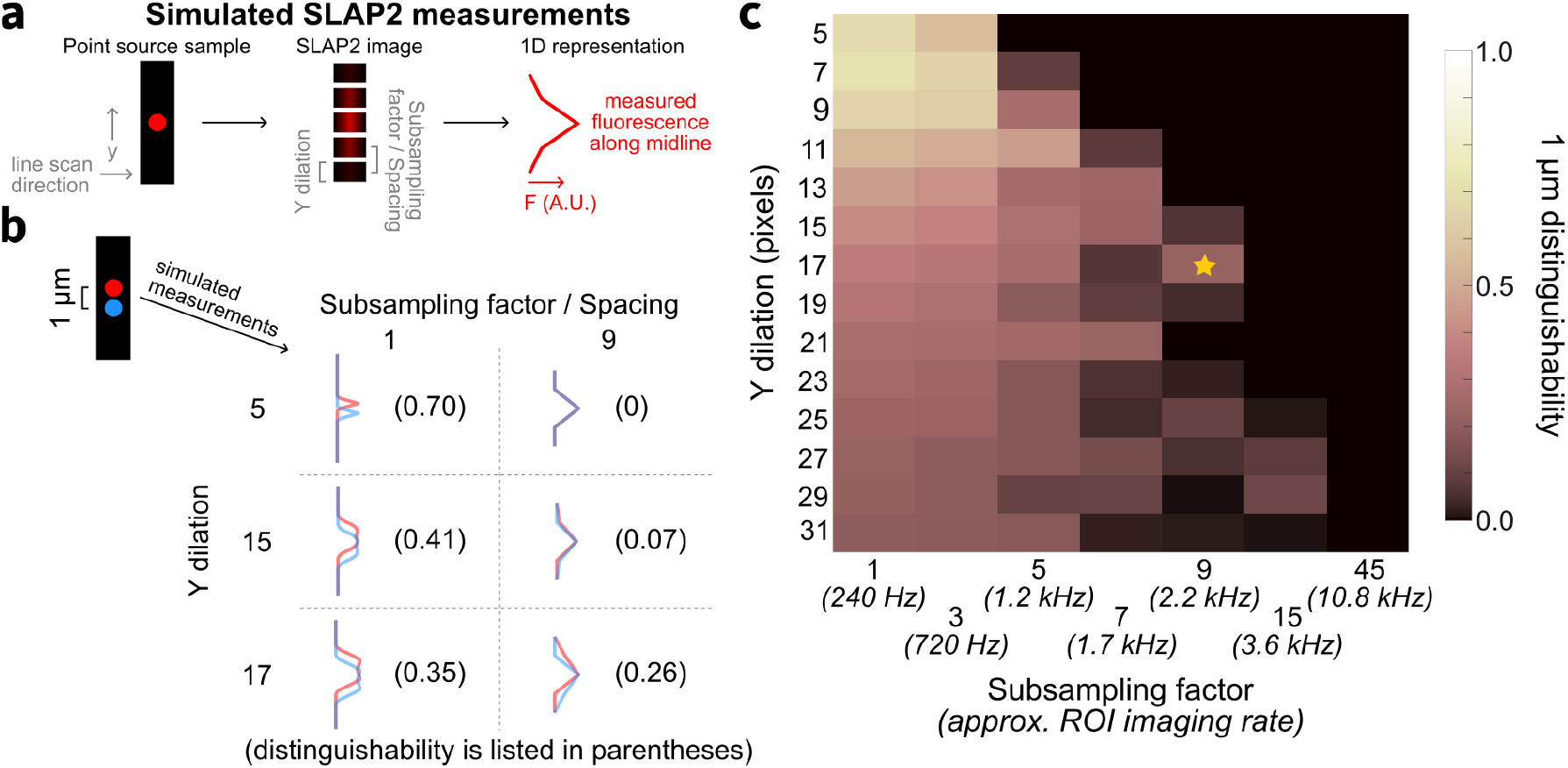
Simulations of SLAP2 band scan parameters, related to Figure 1. **a**, SLAP2 band scan measurements with varied parameter values (Y dilation and subsampling factor) were simulated for point sources and reduced to a 1D representation along the *y* dimension (perpendicular to the line scan direction). **b**, For two point sources spaced 1 µm apart in *y*, the distinguishability (see **Methods**) of SLAP2 measurements was calculated for different combinations of Y dilation and subsampling factor. **c**, The distinguishability of point sources spaced 1 µm apart (1 µm distinguishability) as a function of Y dilation and subsampling factor. To achieve a subsampling factor of 9, the optimal Y dilation of 17 was used (star).

**Figure S5:**
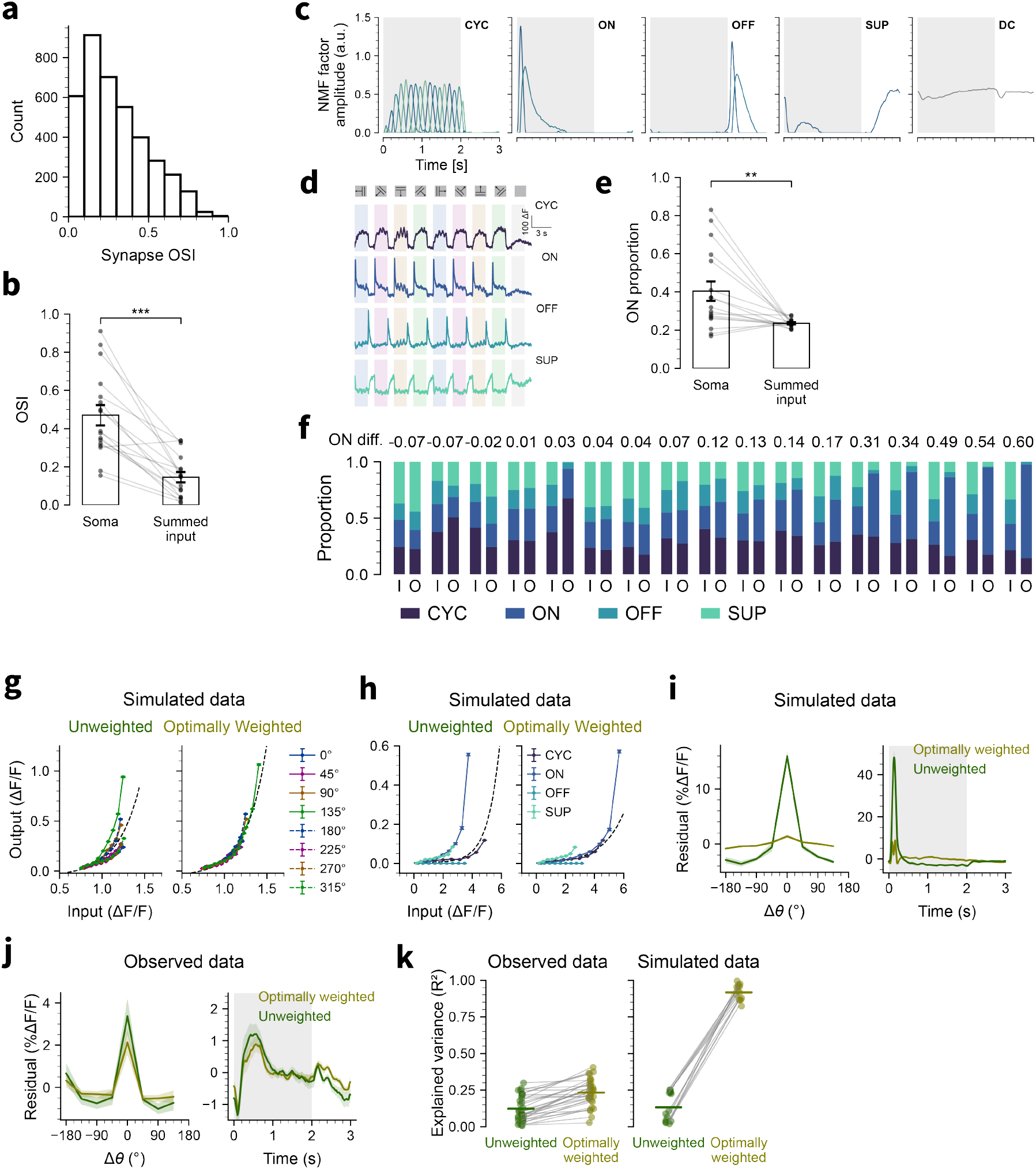
Orientation tuning, temporal factor decomposition and activation functions of synaptic inputs and somatic output, related to Figure 3. **a**, OSI distribution of synaptic inputs. **b**, Paired comparison of OSI between somatic calcium and summed synaptic input (Wilcoxon signed-rank test, *N* = 17 stimulus-modulated sessions). **c**, Temporal factors from NMF decomposition. **d**, Mean tuning profiles of the 100 synapses with the highest loadings in each NMF factor category. **e**, Proportion of the ON temporal factor category in somatic output vs. summed input (Wilcoxon signed-rank test, *N* = 17 sessions). **f**, Proportion of each factor category in the summed input (I) and somatic output (O) for each session (*N* = 17 stimulus-modulated sessions). **g-h**, Same analysis as Figure 3g-h, applied to a simulated neuron whose output was generated by an exponential-nonlinearity GLM (exp(*w ·x*)) of all its synaptic inputs. Only 5% of synapses were observed, and measurement noise was added to the observed synaptic signals and simulated somatic output (see Methods). **i**, Mean residuals from the best GLM fit to simulated datasets (*N* = 10 direction-amplified simulations, left; *N* = 9 ON-amplified simulations, right). “Unweighted” refers to the fit where summed input is the unweighted sum of synapses; “optimally weighted” refers to the fit where each synapse was assigned a single weight from the GLM, and summed input is the weighted sum. **j**, Same as **i**, but for the observed datasets (*N* = 17 sessions). **k**, Explained variance (R2) of the GLM with unweighted and optimally weighted summed input, observed sessions (left, *N* = 17) and simulations (right, *N* = 19) (* *p* < 0.05, ** *p* < 0.01, *** *p* < 0.001)

**Figure S6:**
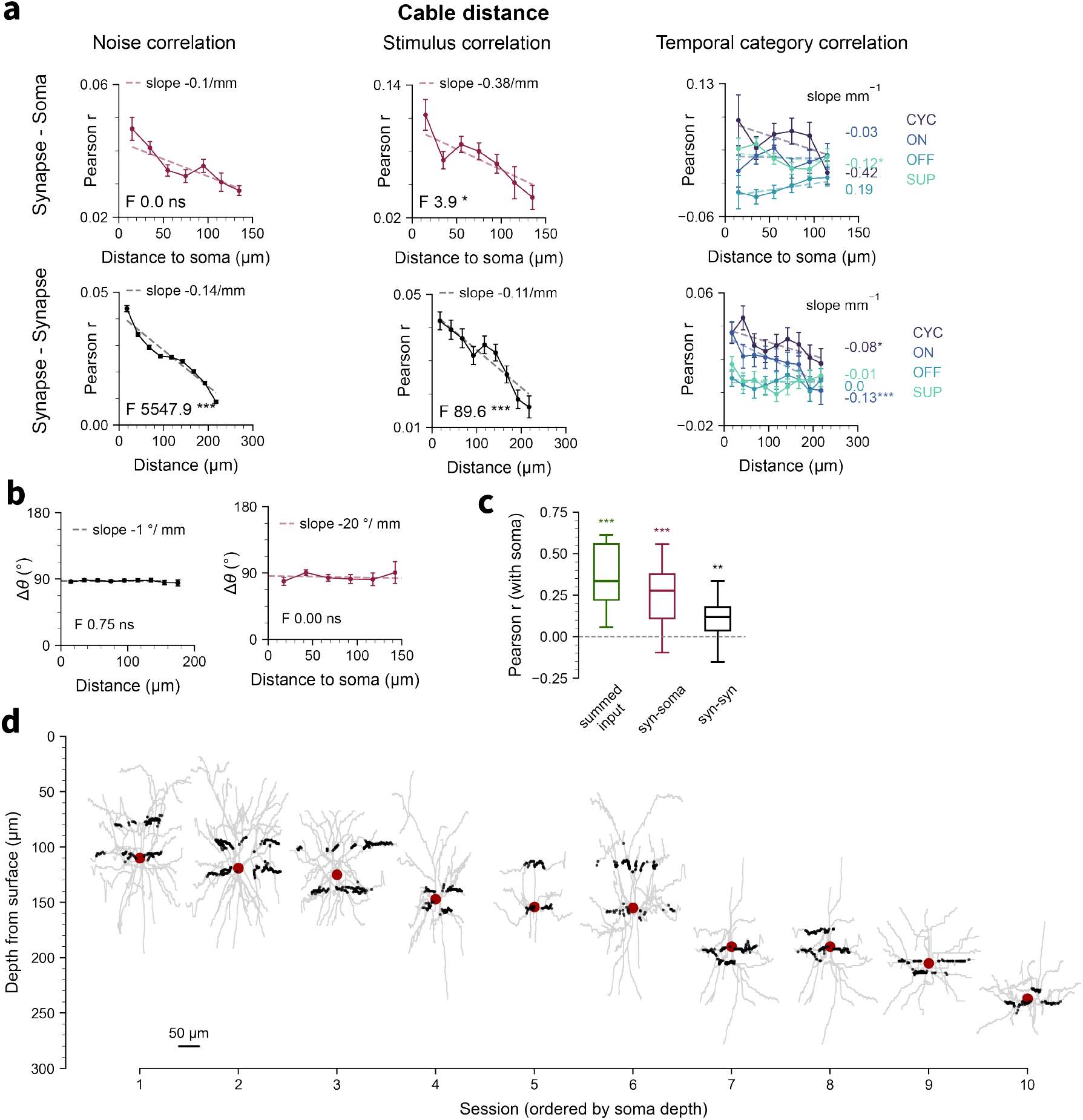
Spatial organization of synaptic tuning and noise correlations relative to the soma, related to Figure 4. **a, left:** noise correlation; **middle:** cross-validated stimulus correlation. **right:** cross-validated stimulus correlation for each temporal factor category, as a function of cable distance between synapses and soma (top) and between pairs of synapses (bottom) (ANCOVA, *N* = 10 sessions). **b**, Difference in preferred direction vs. Euclidean distance for synaptic pairs (left; *N* = 10 sessions), and synapses and soma (right; *N* = 7 stimulus-modulated sessions with soma OSI > 0.3); synapses with OSI > 0.3 (ANCOVA). **c**, Correlation (Pearson’s *r*) between the soma’s tuning profile and the summed input tuning profile (green), mean synapse-soma noise covariance (purple), and mean synapse-synapse noise covariance (black) (one-sample Wilcoxon signed-rank test, *N* = 17 stimulus-modulated sessions). **d**, Depth of soma (dark red) and imaged synapses (black) overlaid on dendritic reconstruction (gray), measured from the pial surface (*N* = 10 sessions, sorted by soma depth). (* *p* < 0.05, ** *p* < 0.01, *** *p* < 0.001)

**Figure S7:**
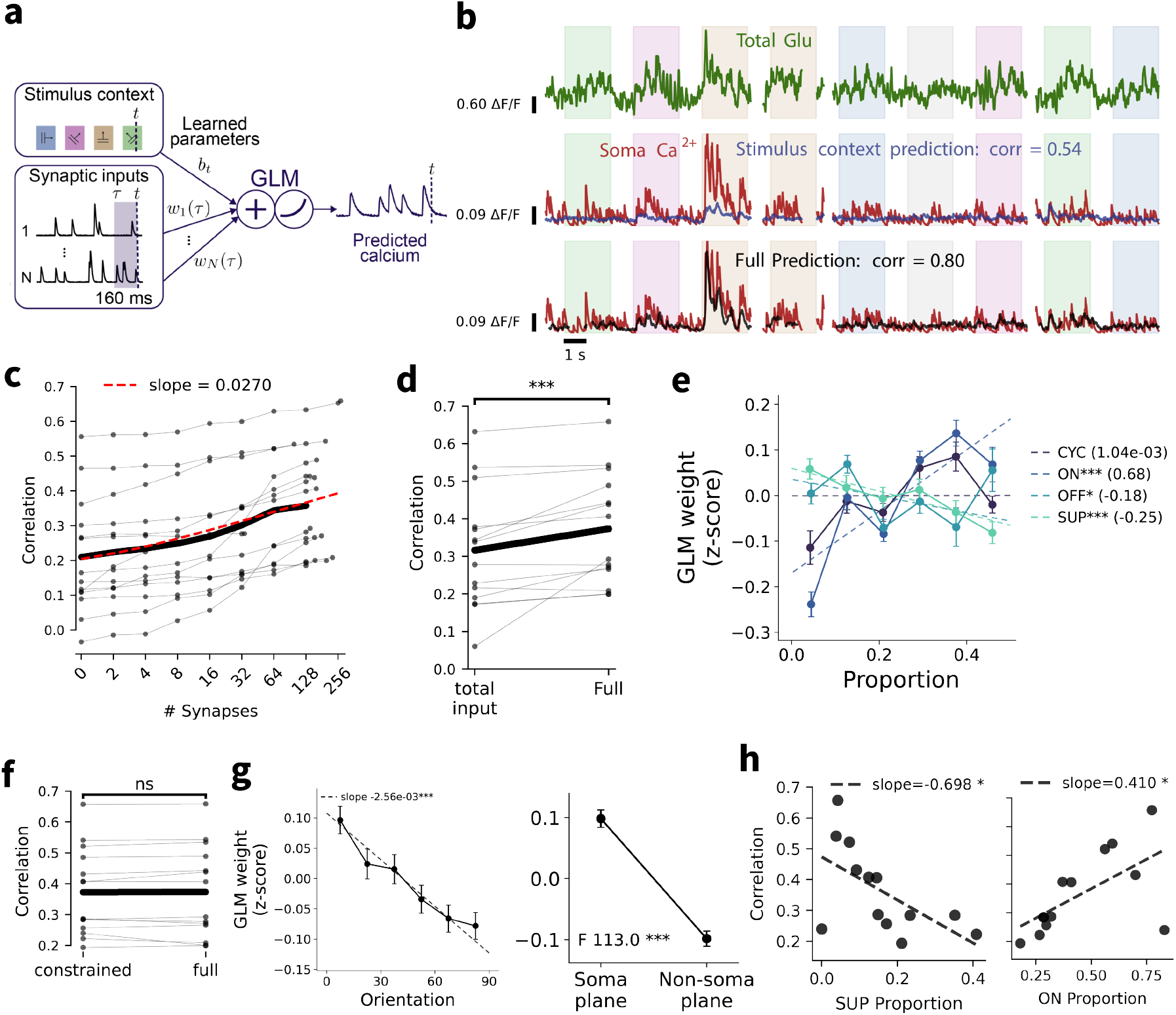
Using generalized linear models (GLMs) to model the input-output functions of neurons. **a**, Schematic of a generalized linear model (GLM) used to predict somatic output from synaptic inputs. Predictions at time *t* are computed from a bias term *b*_*t*_ and the preceding 160 ms of synaptic input. **b**, Example GLM predictions showing summed input (green), true somatic calcium (red), output predicted from only the time-varying bias term *b*_*t*_ (purple) and output predicted from the full model (black). **c**, GLM performance (Pearson *r*) as a function of the number of synapses included in the model. **d**, Comparison of GLM performance (Pearson *r*) between the model that assigns equal weight to all synapses (total input) and the full model, which includes the bias term *b*_*t*_ and assigns a separate weight to each synapse. **e**, GLM weight vs. proportion of each temporal factor category (ON, CYC, OFF, SUP) across synapses. **f**, Comparison of model performance (Pearson *r*) between the full model and constrained model (see **Methods**). **g**, GLM weight vs. preferred orientation relative to soma’s preferred orientation (left) and vs. dendritic plane (right) across synapses. **h**, GLM performance (Pearson *r*) as a function of somatic SUP (left) and ON (right) temporal category proportions. (* *p* < 0.05, ** *p* < 0.01, *** *p* < 0.001)

**Figure S8:**
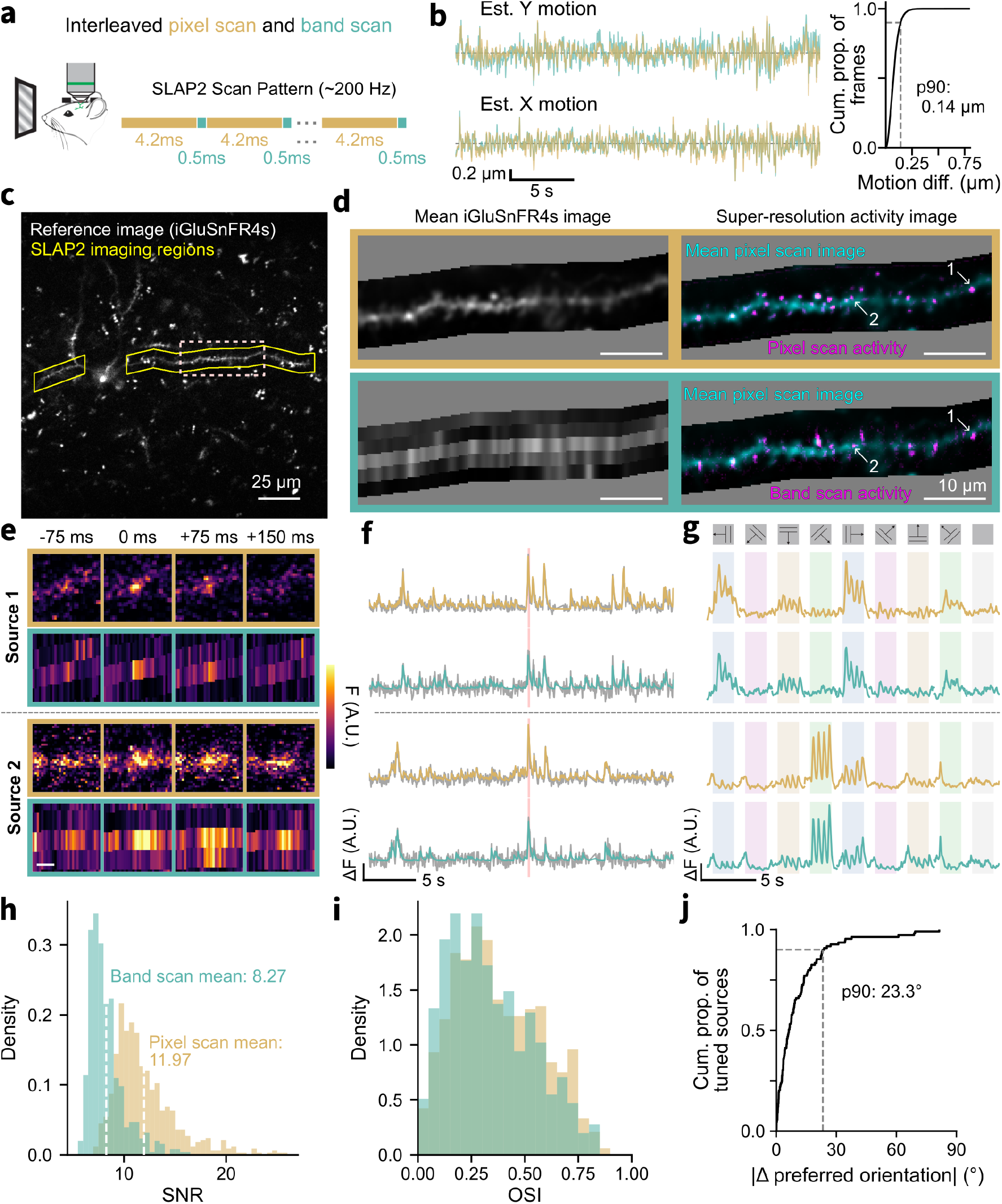
Simultaneous SLAP2 recording in band and pixel scanning modes, related to Figure 5. **a**, SLAP2 pixel scanning and band scanning were interleaved in a single plane SLAP2 imaging experiment while the mouse passively viewed drifting gratings. **b, left:** Example offline estimated motion displacements from pixel scan data (yellow) and band scan data (teal). **right:** Cumulative distribution function of the magnitude of difference in motion displacements between pixel scan and band scan data across all frames of four ~20 minute recordings. **c**, Imaging was performed on select regions of interest (yellow) with iGluSnFR4s-expressing dendrites. **d, left:** Mean iGluSnFR4s fluorescence across the experiment duration for the pixel scan data (top) and band scan data (bottom). The dendrite segment corresponds to the dotted box in (c). **right:** Super-resolution activity image (magenta) from the pixel scan data (top) and from the band scan data (bottom), both overlaid on the mean iGluSnFR4s fluorescence from pixel scan data (cyan). **e**, iGluSnFR4s fluorescence across time around a glutamate transient acquired by pixel scanning (yellow) and band scanning (teal) for the two activity peaks indicated in (d). Scale bar, 2 µm. **f**, Fluorescence traces extracted from the two regions depicted in (e). Grey traces are the least-squares estimated traces. The denoised pixel scan traces and denoised band scan traces are in yellow and teal, respectively. Red highlight indicates the time window shown in (e). **g**, Trial-averaged tuning profiles for the traces shown in (f). **h**, Distribution of peak signal-to-noise ratio (SNR) of extracted pixel scan sources (yellow) and band scan sources (teal). **i**, Distribution of orientation selectivity index (OSI) for sources from pixel scanning (yellow) and band scanning (teal). **j**, Cumulative distribution function of the difference in preferred orientation between strongly orientation selective pixel scan sources (OSI>0.5) and their matched band scan sources.

**Figure S9:**
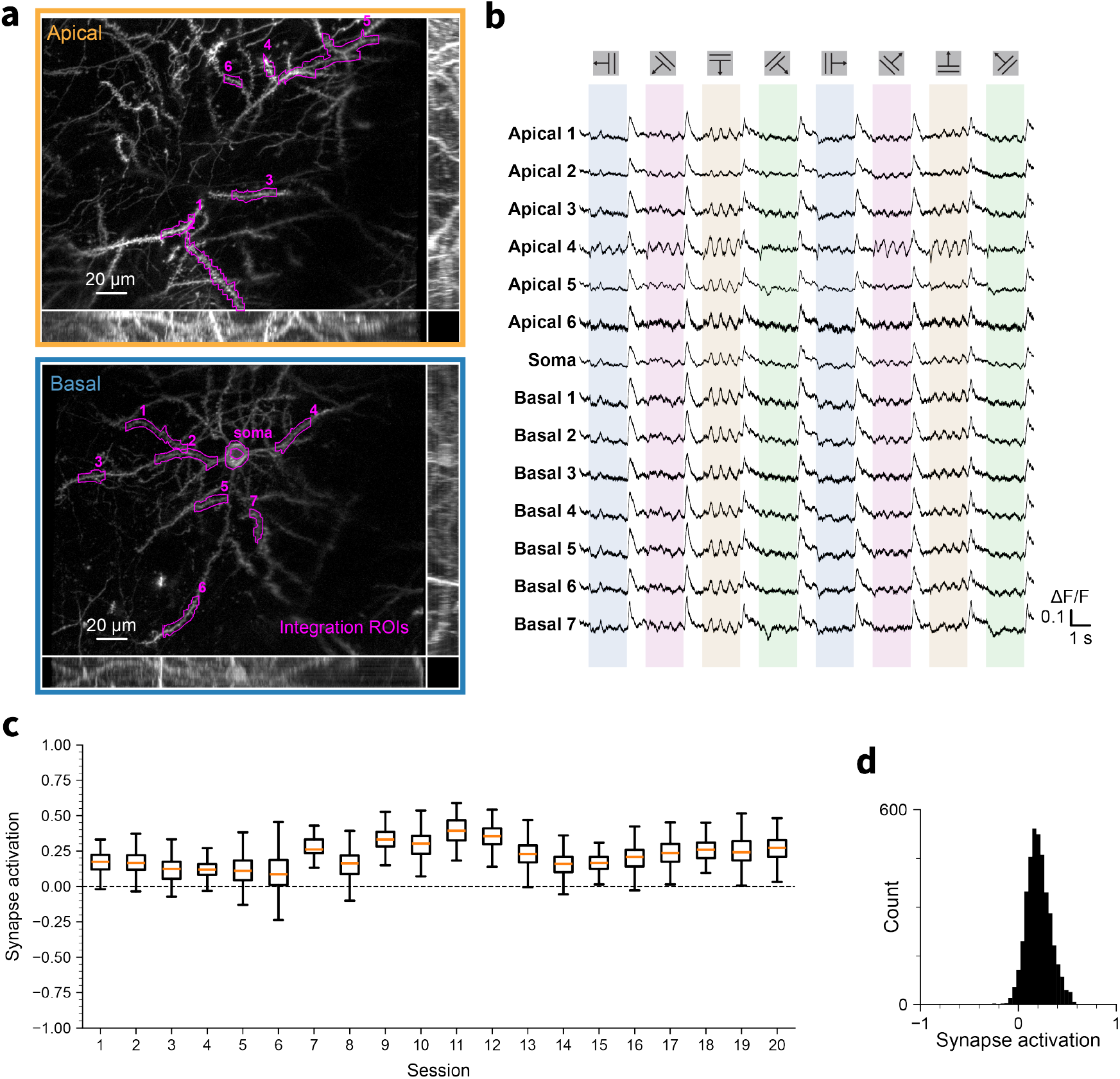
Dendritic voltage responses and synapse activation during oscillatory events, related to Figure 6. **a**, Simultaneous imaging of soma and 13 dendritic segments across basal and apical planes in a layer 2/3 pyramidal neuron. **b**, Trial-averaged stimulus responses of ROIs shown in (a). **c**, Synapse activation during oscillatory events for each session. **d**, Histogram of synapse activation during oscillatory events, across all sessions and all detected oscillatory events.

**Movie S1:**
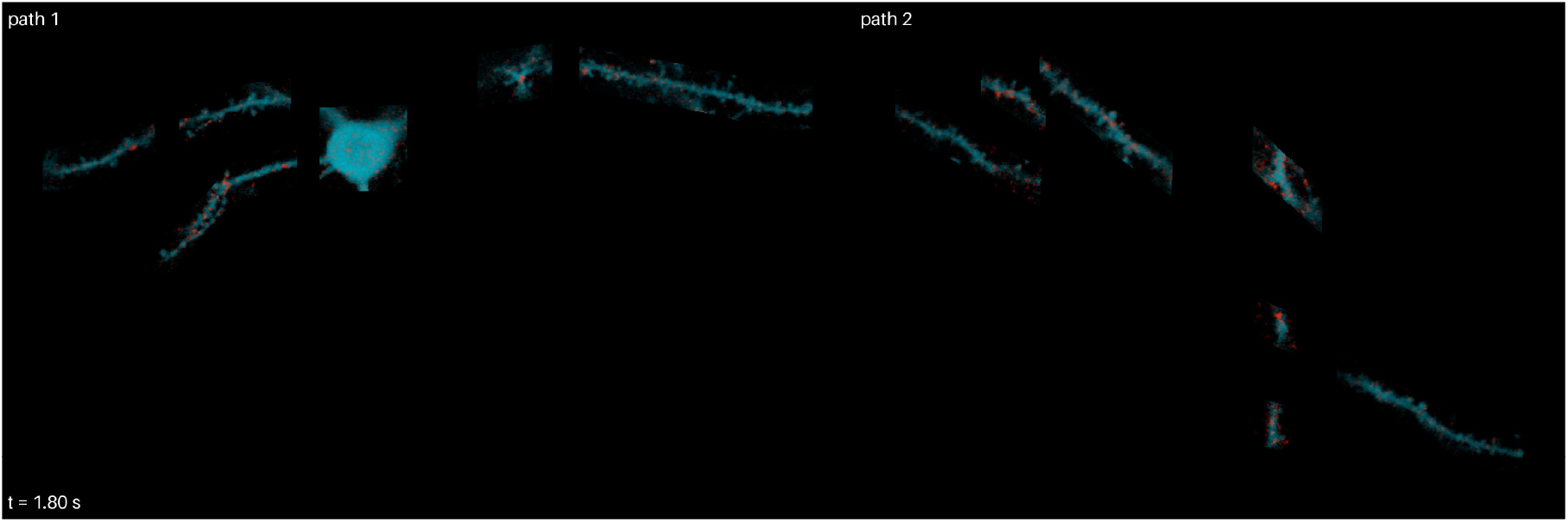
Glutamate imaging with pixel scanning during drifting grating stimulation, related to Figure 2. Single trial of a SLAP2 recording from a layer 2/3 pyramidal neuron, showing the two imaging paths side by side (path 1, left; path 2, right). Mean fluorescence is shown in cyan, overlaid with increases in iGluSnFR4f fluorescence relative to baseline in red (normalized Δ*F*/*F*, saturating at the 99.97th percentile). The square at the bottom left shows the visual stimulus (drifting grating, 2 Hz, 0.04 cycles/degree, 2 s per direction, separated by 1 s of blank screen). Time since trial onset is shown at the bottom left. The movie is played at half real time.

**Movie S2:**
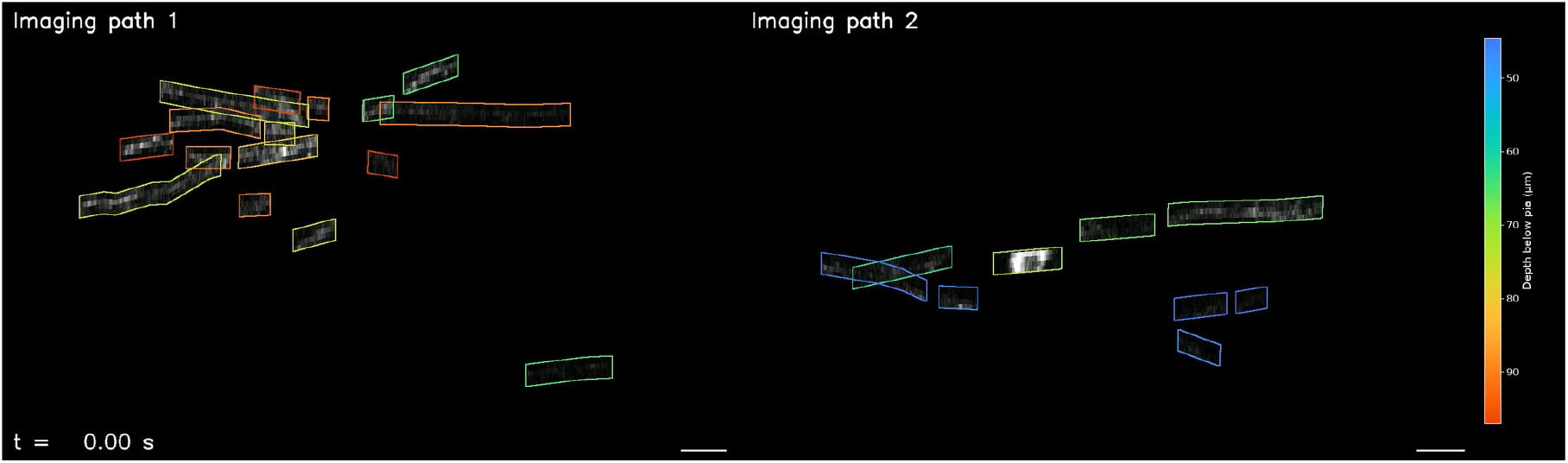
Raw fluorescence movie collected by SLAP2 volumetric band scanning, related to Figure 5. Movie of the fluorescence (maximum intensity projection along depth) captured along dendrites expressing iGluSnFR4s during a segment of SLAP2 volumetric band scanning on the simultaneously active imaging paths. The data were collected at 70 Hz volume rate on path 1 and 92 Hz volume rate on path 2. The first few frames of the movie show a maximum projection intensity of the corresponding reference stack with the boxes indicating the imaging regions (colored by depth from pia) and unimaged regions shaded in a blue tint. Scale bar, 20 µm.

